# Using a barcoded AAV capsid library to select for novel clinically relevant gene therapy vectors

**DOI:** 10.1101/683672

**Authors:** Katja Pekrun, Gustavo De Alencastro, Qing-Jun Luo, Jun Liu, Youngjin Kim, Sean Nygaard, Feorillo Galivo, Feijie Zhang, Ren Song, Matthew R. Tiffany, Jianpeng Xu, Matthias Hebrok, Markus Grompe, Mark A. Kay

## Abstract

While gene transfer using recombinant adeno-associated viral (rAAV) vectors have shown success in some clinical trials, there remain many tissues that are not well transduced. Because of the recent success in reprogramming islet derived cells into functional β-cells in animal models, we constructed two highly complex barcoded replication competent capsid shuffled libraries and selected for high transducing variants on primary human islets. We describe a chimeric capsid (AAV-KP1) that penetrated and transduced primary human islet cells and human embryonic stem cell derived β-cells with up to 10-fold higher efficiency compared to previously studied best in class AAV vectors. Remarkably, this chimeric capsid was also able to transduce both mouse and human hepatocytes at very high levels in a humanized-chimeric mouse model, thus providing a versatile vector which has the potential to be used in both preclinical testing and human clinical trials for both liver-based diseases and diabetes.

## INTRODUCTION

An estimated 30.3 million US Americans are affected by either type 1 or type 2 Diabetes Mellitus (CDC, 2017). Various strategies to cure diabetes have been evaluated over the years with limited success. For example, transplantation of cadaveric human islets into the hepatic duct has been used to replace β-cells in type 1 diabetic patients, so far with low efficiencies (for a review see(Bruni et al, 2014)). Islets are highly vascularized and require large amounts of oxygen to survive. Because re-vascularization of the transplanted islets takes several weeks, those transplants suffer from a large number of cell death due to oxygen deprivation. In order to improve graft survival and function, different approaches of *ex vivo* gene therapy, either by supplying or repressing certain transcription factors using different viral or non-viral delivery systems (for example AAV, adenovirus, lentivirus, various lipids or non-lipid polymers), may be used (see (Wang et al, 2011) for a review). Recently, the transplantation of diabetic mice with islets deficient in PHLDA3, a known suppressor of neuroendocrine tumorigenicity, has been shown to lead to improved glycometabolic condition and improved cell survival during early transplantation (Sakata et al, 2017). Recombinant AAV mediated overexpression of Follistatin, a protein with important roles in the proper functioning of the reproductive, endocrine, and muscoskeletal systems, has been shown to promote β-cell proliferation and maintain pancreatic islet mass in a diabetic mouse model (Zhao et al, 2015). In addition to the approaches described above, it is of utmost importance to prevent loss of the transplanted islets due to recurrent autoimmune destruction. Recently, a study described the use of rAAV to overexpress Igf1 in diabetic mice (Mallol et al, 2017). IgfI is a pro-survival factor and β-cell mitogen that has important roles in β-cell maturation and function and is also involved in the interplay between the endocrine and the immune system. Over a period of 30 weeks, this gene therapy strategy was successfull in counteracting progression to autoimmune diabetes. Another strategy for the treatment of diabetes is the conversion of glucagon producing α-cells or other endocrine or exocrine pancreatic cell types into β-cells by overexpression or repression of certain transcription factors, such as Pdx1, Ngn3, MafA, Pax4, and Arx (Chakravarthy et al, 2017; Collombat et al, 2009; Courtney et al, 2013; Furuyama et al, 2019; Lima et al, 2016; Matsuoka et al, 2017; Wang et al, 2018; Xiao et al, 2018; Zhang et al, 2016) (for a review see (Vieira et al, 2016)). For the studies that utilized rAAV for gene delivery into murine islet cells, AAV8 had been used as this serotype was shown to be efficient for transduction. Recently, intraperitoneal delivery of an AAV8 based vector expressing IL-2 under control of the β-cell type specific insulin promoter has been described to achieve highly specific transduction of β-cells in mice (Flores et al, 2014). Another study found that AAV6 was highly efficient in transducing mouse islets *in vitro* and *in vivo* when using intraductal delivery. However, when delivered systemically, AAV8 proved to be the more robust serotype (Wang et al, 2006). Recently, tyrosine-phenylalanie (Y-F) mutant AAV8 vectors have been reported to achieve up to ten-fold improved gene transfer into mouse islets as compared to wildtype AAV8 (Chen et al, 2017). Interestingly, studies in rats found that AAV5 was the best capsid for islet transduction in this animal model (Craig et al, 2009). AAV2 is the serotype that has been described most frequently for transduction of human islets (Flotte et al, 2001; Kapturczak et al, 2001). More recently, capsid variants AAV-DJ (Grimm et al, 2008) and AAV-LK03 (Lisowski et al, 2014) were found to have better transduction efficiency on human islets as compared to AAV2 and AAV3B, however, a strong preference for α-cells was observed for those capsids (Song, 2017).

Even though different AAV capsids that are capable of transducing human islets have been described, the current low efficiencies limit their broad applicability to diabetes. In view of that, we sought to develop new capsid variants with further enhanced human islet cell transduction efficiency using directed evolution. Directed evolution by DNA shuffling is a powerful tool to mimic natural evolution in a vastly accelerated manner (Stemmer, 1994a; Stemmer, 1994b) and has yielded greatly improved variants in many different areas of research (Apt et al, 2006; Chang et al, 1999; Christians et al, 1999; Crameri et al, 1997; Crameri et al, 1998; Gafvelin et al, 2007; Ness et al, 1999; Pekrun et al, 2002; Powell et al, 2000; Raillard et al, 2001; Soong et al, 2000; Stutzman-Engwall et al, 2005; Wright et al, 2005; Zhang et al, 1997). Most importantly, no *a priori* knowledge about sequence-function relationships is needed, which stands in contrast to rational design strategies. In 2008 our group was the first to report the development of an improved AAV variants by subjecting a capsid shuffled AAV library to multiple rounds of selection on the target cell type (Grimm et al, 2008). Since then, we and others have used this approach to identify AAV variants with improved properties (Asuri et al, 2012; Gray et al, 2010; Grimm et al, 2008; Li et al, 2008; Lisowski et al, 2014; Maguire et al, 2010; Paulk et al, 2018a; Paulk et al, 2018b; Siu et al, 2017; Tervo et al, 2016; Yang et al, 2009; Yang et al, 2011).

In this project, we subjected capsid shuffled barcoded AAV libraries to multiple rounds of selection on primary human islets and analyzed enriched capsid variants for improved transduction efficiency. As previously described by others (Adachi et al, 2014; Davidsson et al, 2016; Herrmann et al, 2018a) the use of barcoded vectors allowed us to follow enrichment of chimeric variants by employing high-throughput sequencing (HTS). Among the candidates tested in our study, three chimeric variants were found to exhibit considerably improved transduction capacity of human islet cells – particularly of β-cells. In addition, these variants showed improved transduction in other non-pancreatic cell types both *in vitro* and *in vivo*. Therefore, these novel capsids may be useful for various gene therapy applications targeting pancreatic islets as well as other tissues relevant for other diseases.

## RESULTS

### Evaluation of parental capsids for human islet transduction efficiency

We first sought to confirm previous data showing that AAV-DJ and AAV-LK03 have higher human islet transduction efficiency than the closely related natural serotypes AAV2 and AAV3B(Song, 2017). We transduced human islets with GFP expressing rAAV vectors packaged with the different capsids and measured transduction efficiency using flow cytometry analysis (Figure 1A). Due to the limited availability of patient derived islets we did not run replicates in this and several other experiments in this study. As expected, AAV-DJ and AAV-LK03 had higher rates of transduction in human islets than AAV2 and AAV3B.

**Figure 1:**
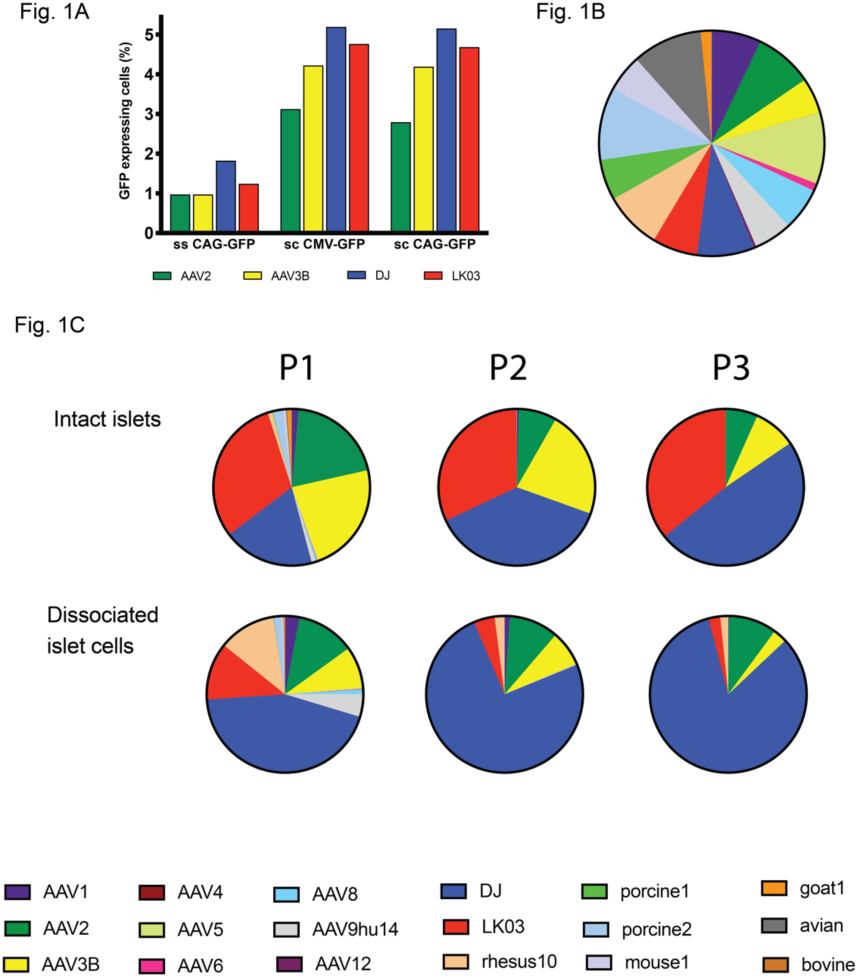
Evaluation of parental capsids for islet tropism. **(A)** Dissociated islet cells were transduced with different vectors that had been packaged with AAV2, AAV3B, DJ and LK03 capsids at a MOI of 1K and transduction efficiency was determined by flow cytometry 48 hrs post transduction. **(B)** Parental contribution in the 18 parent pool as analyzed using high-throughput sequencing of the barcodes. **(C)** Barcode sequences were amplified from viral genomes after passaging the 18 parent pool on human islets and were analyzed by high-throughput sequencing. Color coding of enriched parental capsids for passaging of the 18 parent mix is identical to that of the parental pool.

This confimatory result provided the basis for a validation study. For this experiment, we generated a control AAV pool containing parental AAVs that each carried a unique barcode (BC) sequence immediately downstream of the capsid coding sequence. After mixing equal copy numbers of each of the 18 parental AAVs, we purified the pool and analyzed its composition by HTS of the barcodes (Figure 1B, Table S1). Although some of the parental AAVs were under- or over-represented in the pool, the 18 parent pool was found to be useful in a pilot library screen as a first validation for the use of barcode sequencing in a capsid selection screen. To evaluate which capsid was best at transducing human islets, we infected human intact as well as dissociated islets with the 18 parent pool and super-infected with human adenovirus 5 (Ad5) helpervirus to replicate AAVs prior to infection of the next selection round. Helpervirus-mediated replication ensured that only those AAV variants that had successfully undergone all steps involved in transduction of the target cell would be available for infection of the subsequent round. Moreover, this amplification of enriched AAVs greatly facilitated rescue of the capsid sequences by PCR. We used a ten-fold higher multiplicity of infection (MOI) for infection of intact (20K) than for dissociated (2K) islets since we reasoned that cells located in the center of islets would be harder to infect than those on the periphery. Three consecutive rounds of selection were performed and AAV replication (Figure S1) as well as composition of the viral pool (Figure 1C) was assessed at each round. For rounds 1 and 2 of selection, lower copy numbers of viral genomes were recovered after propagation on intact islets as compared to the input whereas replication reached the highest levels at round 3. When a lower MOI of 2K was used to infect dissociated islet cells, the amount of virus obtained after islet culture was higher than the input at each round of selection. HTS of the BC sequences was used to assess the composition of AAVs at each round. Between 400,000 and 1,000,000 sequence reads were obtained for each sample. Certain parental AAVs had a clear advantage over others as the diversity of the pool was already diminished after one round of selection, particularly when intact islets were used (Figure 1C). AAV2, AAV3B, as well as shuffled variants AAV-DJ and AAV-LK03 were enriched already after the first passage. After the third passage on intact islets, as expected, AAV-DJ and AAV-LK03 were equally represented while AAV2 and AAV3B were present at lower concentrations. Passaging of the 18 parent pool on dissociated islet cells revealed early enrichment of AAV-DJ with lower proportions of AAV2, LK03, AAV3B, and AAV-rhesus10. This experiment validated our selection screen approach as it confirmed the results from previous transduction studies showing that AAV-DJ and AAV-LK03 transduce primary human islets better than closely related serotypes AAV2 and AAV3B.

### Characterization of highly diverse barcoded capsid shuffled AAV libraries

We first generated a library with random barcodes (BC) downstream of the cap polyadenylation site. Insertion of the BC sequence did not negatively impact virus function as shown by replication studies for wild-type AAV with and without the BC (data not shown). HTS of the barcode plasmid library showed a high degree of diversity with over 93% of all BC reads containing a unique sequence (Table S1). As HTS of the barcodes involved an amplification step, it is most likely that the actual number of repetitive barcodes in the BC library was lower than that. The size of the barcode library was estimated to be 1×10^7^ based on the number of colonies obtained from an aliquot of the transformation reaction (data not shown). Two shuffled capsid libraries were then generated using sequences from 10 related AAVs (10 parent library) or 18 more diverse AAVs (18 parent library) and cloned into the BC library vector. A phylogenetic analysis of the parental sequences used for shuffling is depicted in Figure S2A. Importantly, two of those capsid sequences – DJ and LK03 – had been derived from previous screens performed in our laboratory using earlier generation libraries and were added as parentals. The capsid shuffled barcoded 10 parent library and the 18 parent library each each had a size of approximately 5×10^6^ clones. Given the low number of replicates after BC sequencing the complexity was estimated to be at least 1×10^6^ for each of the libraries.

The degree of library diversity was assessed using three different methods: HTS of the barcodes (MiSeq platform), PacBio sequencing of the capsids including the barcodes, as well as Sanger sequencing of the capsids including the barcodes. Amplification and HTS of the barcodes from the plasmid libraries as well as the AAV libraries revealed a very high degree of complexity with low numbers of replicates. At least 1 million reads were obtained for each sample (Table S2). When the 10 parent library was analyzed on the plasmid level, i.e. prior to production in 293T cells, it was found that over 88% of all reads contained unique sequences, with a maximum number of replicates found to be 7 out of 1.25 million reads. For the 18 parent plasmid library, more than 88% of the reads were unique -the maximum number of reads obtained for one single BC sequence was 8 out of 1.8 million reads. Analysis of the NGS data for the AAV libraries revealed approximately 76% and 80% unique BC sequences for the 10 parent and 18 parent libraries, respectively (Table S2).

Several capsids from the 10 parent and the 18 parent plasmid libraries as well as from the 10 parent AAV library were also analyzed by Sanger sequencing and subjected to recombination analysis using the Xover program (Huang et al, 2016) (Figure S3A, Figure S4A). Since parental capsids DJ and LK03 are chimeras derived mostly from serotypes also used in this study (Figure S2B), they are not included as separate parental sequences here. For analysis of diversity on the AAV level, the region spanning capsid and barcodes was amplified from extracted viral genomes, cloned and sequenced (Figure S3A, right panel). In addition to depicting crossover figures of the shuffled capsids, we used the parental contribution data generated using the Xover software to perform conservation analysis throughout the capsid sequences. The conservation values for each amino acid residue were calculated (Figure S3B, Figure S4B) allowing us to evaluate if sequence diversity within the libraries was drastically diminished when compared to the parental sequences. The conservation patterns of the10 parent library on the plasmid as well as the AAV level closely matched those of the parental sequences (Figure S3B). On the other hand, the 18 parent plasmid library exhibited higher conservation levels than the parents throughout most of the capsid (Figure S4B). Crossover analysis of the capsid amino acid sequences showed that the 10 parent library was well shuffled with an average rate of 11 crossover events per capsid. With an average of 8 crossovers, the 18 parent library was less thoroughly shuffled. This observation can be explained with the large proportion of parental capsids that have low homology to each other throughout large stretches. Crossovers based on an identity of less than 15 bases are difficult to obtain (Stemmer, 1994a). Thus it is not surprising that fragments with low homology to each other do not easily recombine when equimolar amounts of each parent are mixed together. The observation that fewer crossovers were found within the 3’ half of the library capsids can also be explained by this limitation as this part of the capsid contains several stretches of low sequence homology between the different serotypes. Complexity of the 10 parent library on the AAV level was found to be similar to that on the plasmid level indicating that capsid assembly during AAV production in 293T cells does not reduce diversity (Figure S3). Moreover, the crossover analysis patterns show that the parent serotypes appear to be represented roughly equally and randomly within the libraries, although AAV3B sequences seem to be slightly favoured in the 3’ part of the shuffled clones. In addition to Sanger sequencing of individual capsids, we performed PacBio sequencing of the 10 parent library on the plasmid level prior to transfection into 293T cells (Figure 2A) as well as on the AAV level (Figure 2B). We obtained 21,000 full-length reads for the 10 parent plasmid library and about half of that read number for the 10 parent AAV library. The graphs in Figure 2 depict the contribution of each parental amino acid residue throughout the length of the capsid. As already observed by Sanger sequencing, PacBio sequencing revealed that the 5’ half of the capsid gene contained a more even distribution of all parental sequences than the 3’ half. Particularly residue contributions from AAV-porcine2 and AAV9hu14 were starkly diminished in the sequence stretch between amino acid positions 450 and 650 while AAV3B sequences were overrepresented. An overrepresentation of AAV3B sequences is possibly at least in part due to the fact that LK03, which contains large sequence stretches from AAV3B (Figure S2B), was used as a parental for shuffling.

**Figure 2:**
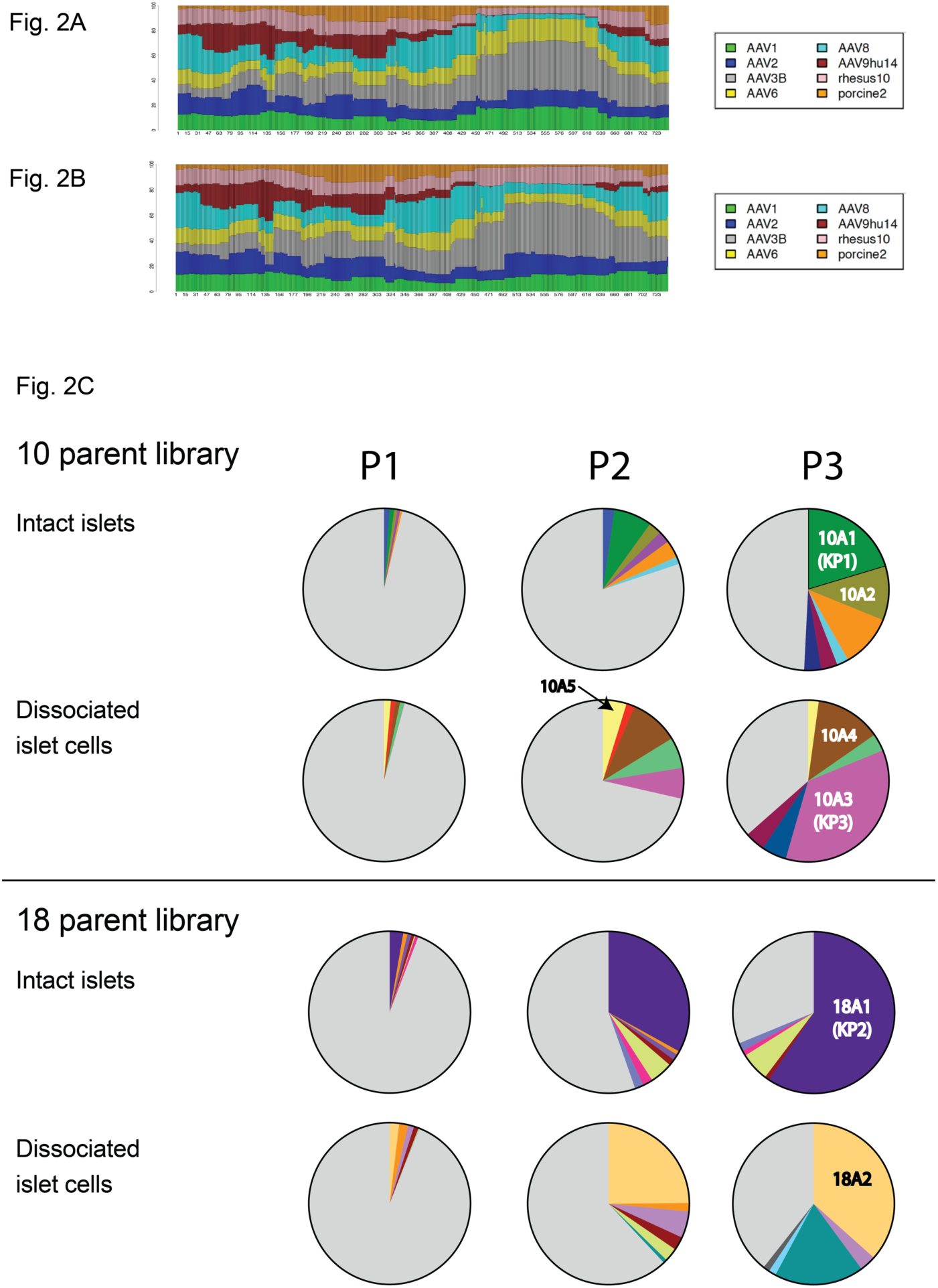
Parental contribution of the 10 parent library and enrichment of distinct capsids during passaging. Chimeric capsid sequences obtained from the 10 parent library on the plasmid level **(A)** and on the AAV level **(B)** were analyzed by PacBio sequencing and parental contribution was assigned for each amino acid position. **(C)** Barcode sequences were amplified from viral genomes after each passage and were analyzed by high-throughput sequencing. Enriched variants are depicted in different colors while all other variants are shown in grey. Enrichment of AAV capsid variants used for vectorization is indicated (10A1, 10A2, 10A3, 10A4 10A5, 18A1, 18A2).

We also analyzed if each capsid was linked to a unique barcode sequence and found very high levels of capsid-BC linkage. According to PacBio sequencing, only 0.5% of all sequences had different capsids sharing identical barcodes.

### Selection of AAV capsids with human islet tropism

Both libraries were used to infect intact as well as dissociated islets and were replicated using Ad5 superinfection. After each round of selection, virus replication was assessed by qPCR (Figure S1) and enrichment of variants was tracked by HTS of the barcodes (Figure 2C). The number of reads was between 400,000 and 800,000. In contrast to the 18 parent pool we observed replication of the libraries at each round of selection which might indicate that certain capsid sequences conferred improved tropism, transduction, and/or replication. Enrichment of distinct capsid variants was observed early in the selection process and are depicted as colored slices in the pie charts in Figure 2C. A second set of library screen on intact islets using two different MOIs was performed in duplicates for the 10 parent library and in triplicates for the 18 parent library. Replication data for this library screen are shown in Figure S5A, the BC sequencing data are shown in Figure S5B. We observed similar levels of replication and enrichment during this second set of library screens. However, this time one BC was found to be enriched in two independent screens of the 10 parent library (variant 10B5, highlighted in yellow), indicating that this particular variant may have a significant selection advantage in islets. A different BC was found to be enriched in two of the triplicate screens of the 18 parent library (18B2, highlighted in magenta).

### Recovery of capsid sequences and evaluation of selected AAV capsid variants for improved transduction of primary human islets and β-cells derived from human embryonic stem cells

A total of 17 enriched capsid sequences were recovered from the different screens after three rounds of selection (see Figure 2C and Figure S5B for enrichment data of those variants) using a forward primer upstream of *cap* and a BC sequence specific reverse primer (Figure 3A), followed by TOPO cloning, and sequencing. After performing sequencing analysis, it was found that all enriched capsids shared large sequences from AAV3B in the 3’ half while the 5’ half was much more diverse (data not shown). All 17 capsid variants as well as one of the control capsids (AAV-LK03) were used to package an AAV vector containing a CAG-GFP cassette. Three of the capsids (18A2, 10B2, and 10B4) failed to generate high-titer rAAV and were excluded from further testing. In an initial pre-screen, crude cell lysate derived rAAV was used to transduce dissociated islet cells using a low MOI. Transduction efficiency was determined by flow cytometry analysis of GFP expressing cells two days post transduction. While multiple capsid variants exhibited higher transduction efficiency than AAV-LK03, others were only marginally improved or not improved at all (Figure 3B). The three lead candidates (10A1, 18A1, and 10A3) were all derived from the first set of library screening and were re-named into KP1, KP2, and KP3, respectively. As we had established in earlier experiments that both AAV-DJ and AAV-LK03 capsids transduced islets with higher efficiency than AAV2 or AAV3B (Figure 1A), only these two capsids were used to perform comparative studies with the novel selected AAV capsids variants. Dissociated islet cells were transduced with purified capsid variants as well as with AAV-DJ and AAV-LK03 at three different MOIs and transduction efficiency was analyzed by flow cytometry. As it was detected before, these three AAV variants were capable of transducing islet cells with improved efficiency when compared to the best parents (Figure 3C, Figure S6A). In fact, these levels of transduction were achieved by AAV-DJ or AAV-LK03 only when a 10-fold higher MOI was used. Next, we wanted to determine if the novel capsids transduced both α- and β-cells with equal efficiency, or if one cell population was being targeted preferentially. Due to limited islet availability only two of the novel and apparently most efficient AAV capsids were used in the study. In order to address this, islets were transduced with GFP expressing vectors packaged into KP1, KP2, DJ and LK03 capsids and the different subpopulations were separated using specific antibody staining for α- and β-cells. KP1 and KP2 remarbably outperformed DJ and LK03 in β-cells but only modestly in α-cells (Figure 3D). Importantly, these data also demonstrate that the novel AAV variants were capable of penetrating intact islets and transduce almost all of the α- and β-cells when using high MOIs.

**Figure 3:**
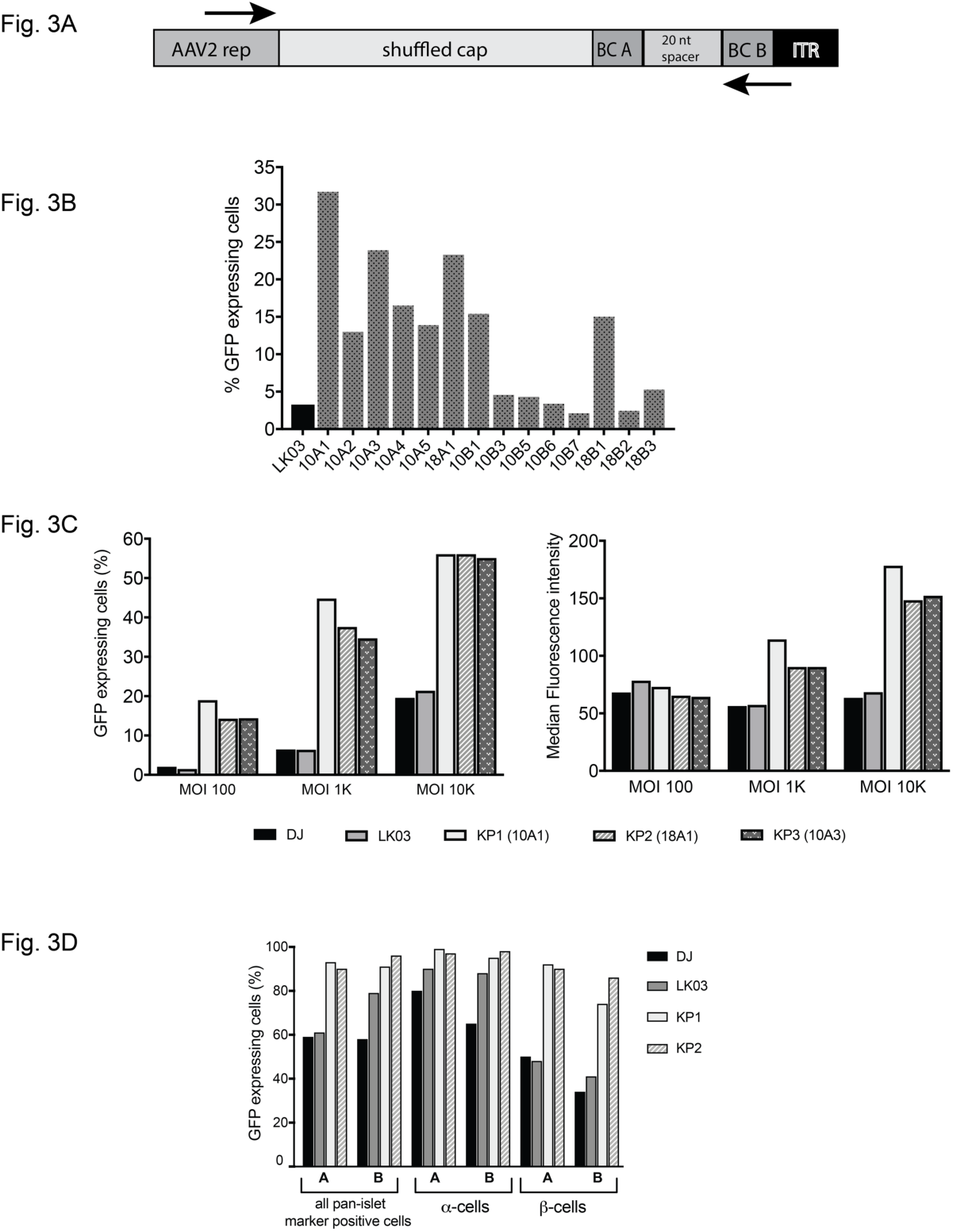
Rescue of enriched capsid sequences and evaluation of selected capsids for islet transduction. **(A)** The forward primer annealed to a sequence in the 3’ end of the rep gene, the reverse primer was specific to the sequence of the right barcode of the variant capsid to be amplified. **(B)** A self-complementary AAV expressing GFP was packaged with LK03 as well as 12 novel capsid sequences and islet cells were transduced using a low MOI of 1,000. Cells were sorted for GFP expression using FACS 48 hrs later. **(C)** Dissociated islet cells were transduced with CsCl gradient purified scCAG-GFP rAAV preps generated with the two best parental capsids as well as the novel capsids that were the top transducers in the pre-screen. Three different MOIs were used for transduction. Transduction efficiency is depicted both as the percentage of GFP positive cells (left graph) as well as the median fluorescence intensity within the GFP positive cell population (right graph). **(D)** α- and β-cell specific transduction efficiency of the novel variants. GFP expressing rAAV packaged with two of the novel variant capsids as well as AAV-DJ and AAV-LK03 capsids were used to transduce intact islets from two individual donors (A, B) at a MOI of 10,000 and α- and β-cell specific transduction was determined by surface staining followed by flow cytometry.

Beta-cells derived from human embryonic stem cells (hESC) are currently being studied as a potential cell source for the treatment of patients with diabetes. Thus, we decided to test if our variants would also transduce these cells with high efficiency. Due to the limited availability of hESC and the labor intensity of the protocol for β-cell generation that we had recently developed(Nair et al, 2019), we focused on the most powerful capsid variant for this study - KP1. Tomato Red expressing vectors were used in this experiment and cells were stained for the β-cell marker C-peptide (Figure 4, Figure S6B, Figure S6C). Similar to what had been observed in islet cells, transduction of these cells using AAV-KP1 was 5- to 10-fold more efficient as compared to AAV-DJ or AAV-LK03.

**Figure 4:**
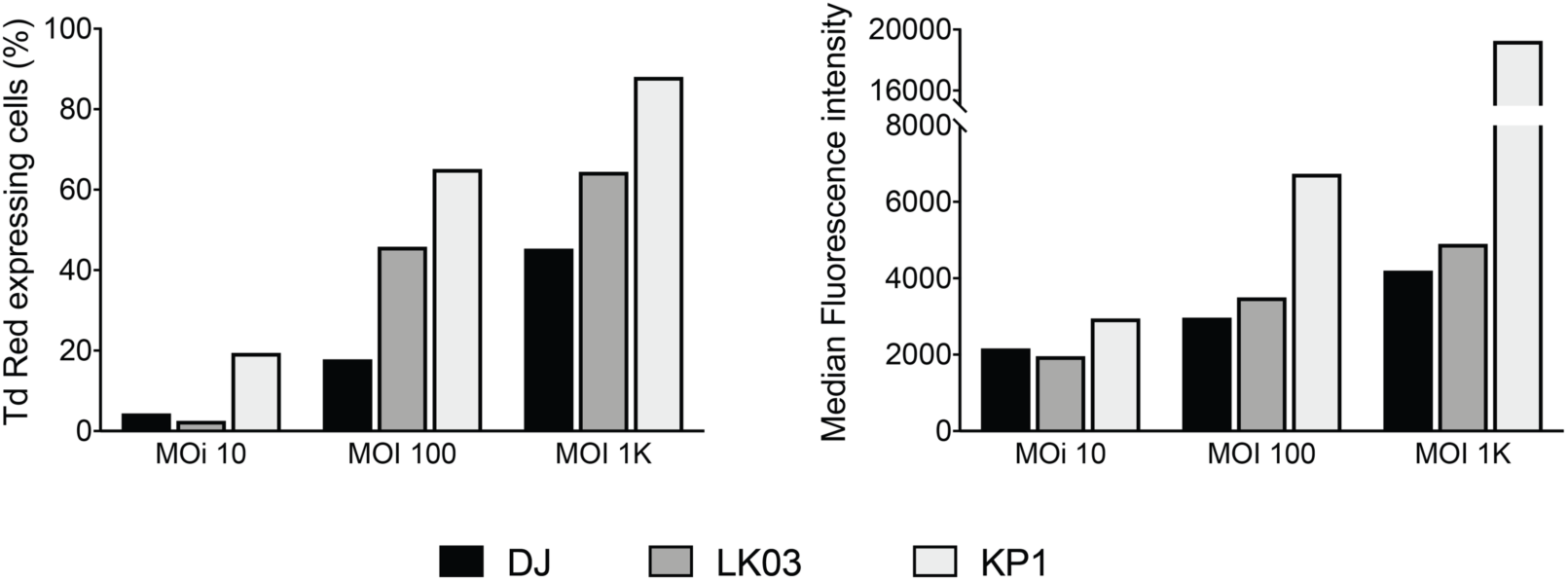
Transduction efficiency of capsid KP1 for human embryonic stem cell derived β-cells. DJ, LK03 and KP1 capsids were used to package a Tomato Red vector and hESC derived mature β-cells were transduced with the MOIs indicated. Intracellular staining for the β-cell marker C-peptide was performed on day 6 post transduction and cells were analyzed by flow cytometry.

### Analysis of sequence determinants of the novel capsids

In Figure 5A the crossover analyses of the three most improved capsid amino acid capsid sequences are shown. While the N-terminal part of capsid KP1 contains stretches from several different parental sequences, capsids KP2 and KP3 are less diverse in this part. However, as seen for all capsids enriched during the islet screen, most of the C-terminal half of those capsids was derived from AAV3B, suggesting that this stretch may contain sequence determinants that are crucial for islet cell tropism. AAV3B is the parental that is most closely related to the three novel capsids with 92% sequence identity to KP1 and KP3 and 95% identity to KP2 throughout the entire capsid sequence. According to analysis of the nucleotide sequences (Figure S7), the KP1 capsid sequence contains fragments from at least seven out of the eight parental serotypes, while KP2 and KP3 contain fragments from at least six parental capsids. The parental contributions were very similar to those found for the amino acid sequences for most part, however some parental contribution patterns differed. As an example, in the case of KP2 capsid the region around nucleotide position 900 was derived from AAV-porcine2 capsid rather than AAV1 or AAV6 capsid as the amino acid crossover analysis had suggested. The Xover program will always attempt to minimize the number of crossover events. Since all the parental sequences share a common amino acid sequence between amino acid position 275 and 312 (Figure S8A), but have different nucleotide sequences, this stretch is shown as being derived from AAV1 or AAV6 when using amino acid sequences, but is shown as being derived from AAV6 and AAV-porcine2 when performing the analysis using the nucleotide sequences. The fact that all three improved capsids share three residues that are unique to AAV1 and AAV6 in the sequence stretch between aa 225 and 267 may indicate that these residues are important for human islet transduction (Figure 5A, Figure S8A). Since AAP uses a different reading frame within the capsid gene, we also performed crossover analysis for this protein. We found that all three novel variants contained chimeric AAP sequences (Figure 5B, Figure S8B). As seen for the nucleotide sequence crossover analysis several KP2 AAP residues were derived from AAV-porcine2. Besides the crossover analysis, we performed enrichment analysis that confirmed strong selection pressure for certain amino acid residues in all three capsids (Figure 5C). In the N-terminus, the two most improved variants KP1 and KP3 showed a strong selection of AAV2 residues while they share AAV8 derived residues in the C-terminus. KP2 capsid shows a strong enrichment of several AAV2 residues between positions 150 and 210. All three capsids have an arginine at the position that has been described to be the key HSPG binding site for AAV3B (position 597 in Figure S8A) (Lerch & Chapman, 2012). Capsids from clade C AAVs that were originally isolated from human specimen (Gao et al, 2004) have been found to be recombinants between sequences closely related to AAV2 (N-terminal half) and AAV3B (C-terminal half). We performed phylogenetic analysis of our novel capsid variants with the 8 parental sequences as well as several members of clade C (Figure S9a) and found that KP1, KP2, and KP3 are more closely related to AAV3B than the clade C capsids. We also ran crossover analysis of clade C as well as the novel capsids using only AAV2 and AAV3B as parental sequences and while both groups share similar patterns overall our novel capsids are less similar to AAV2 in the N-terminal half and are more similar to AAV3B in the C-terminal part than clade C capsids (Figure S9b). We also performed predictive three-dimensional structural VP3 capsid mapping of the novel variants to reveal those residues that are displayed on the outside of the capsids and thus may contribute to improved islet transduction, possibly by interaction with cell surface receptors (Figure 5D). All amino acid residues that differ from the closest parental AAV3B and are shared among at least two of the novel variants are listed in Table S2. The three novel capsids also share several non-AAV3B derived residues within VP3 that are buried inside the capsid as well as residues in the N-terminus of the capsid gene which encodes parts of VP1 and VP2 (Table S3). Those residues possibly also contribute to the observed strong transduction efficiency as they may be involved in uncoating and other post entry steps. All but one of the surface-exposed tyrosine residues that have been associated with ubiquitination and subsequent proteasome-mediated degradation of AAV2 capsids are present in the novel variants (Zhong et al, 2008). However, all variants have a phenylalanine at one of the positions where AAV2 contains a tyrosine since this stretch is derived from AAV3B (position 504 in Figure S8A).

**Figure 5:**
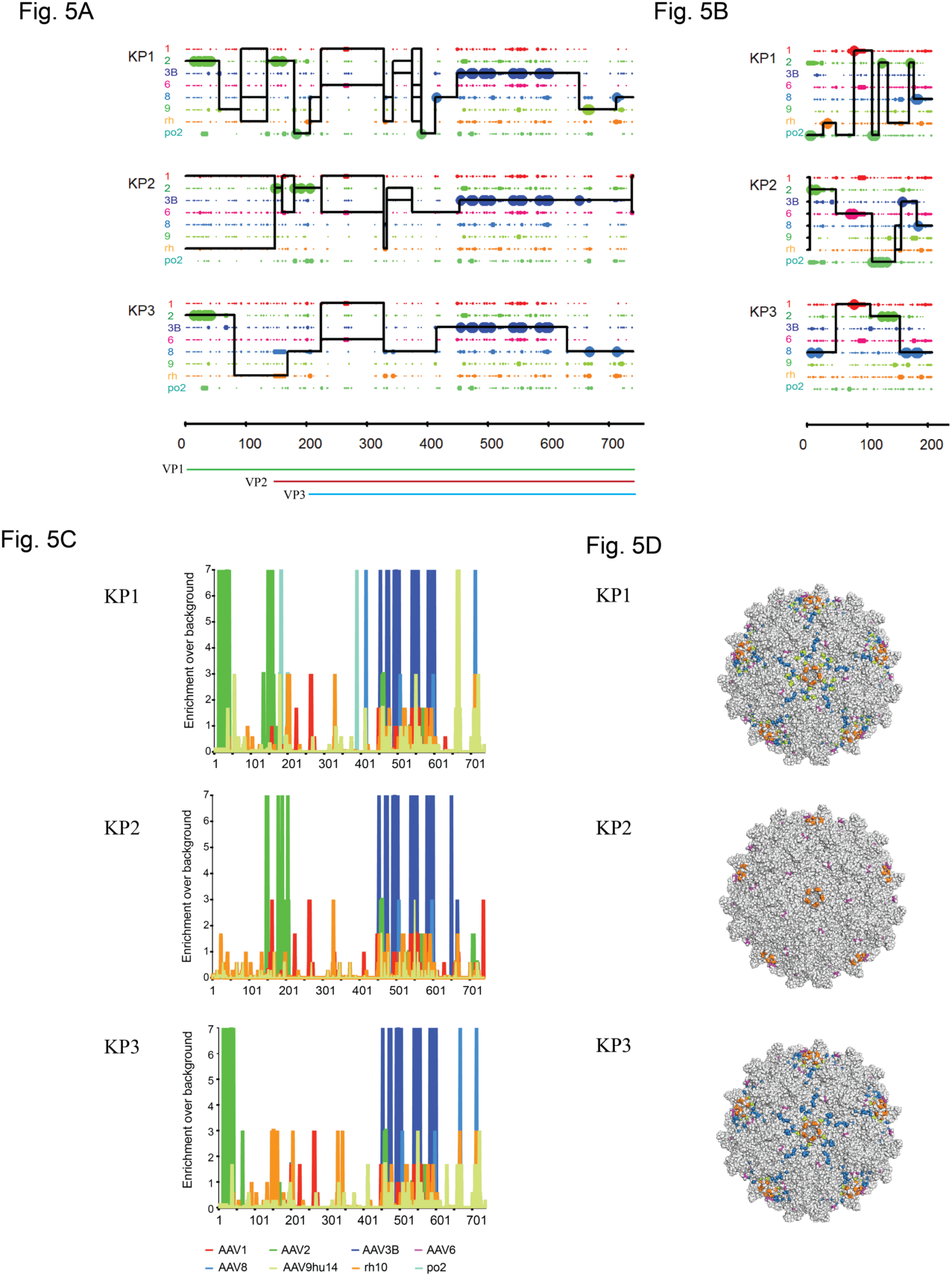
Amino acid sequence and structural composition of selected shuffled AAV capsid variants. (**A**) Amino acid sequence mapping analysis of parental capsid fragment crossovers in vectorized shuffled capsids. Library parents are depicted in different colors as indicated on the left. Large dots represent 100% parental match (i.e. the position in question matches only one parent) and small dots represent more than one parental match (i.e. the position matches more than one parent) at each position. The solid line for each chimera represents the library parents identified within the sequence between crossovers. A set of thin horizontal parallel lines between crossovers indicates multiple parents match at an equal probability. A vertical spike indicates a single position switch between parents. VP1, VP2, VP3 and AAP ORFs are shown below. **(B)** Amino acid sequence mapping analysis of parental AAP fragment crossovers in vectorized shuffled capsids. **(C)** Enrichment scores were calculated for each amino acid position in the sequence of each chimera by comparison of sequences from parental serotypes based on maximum likelihood. Library parents are depicted in different colors as shown. **(D**) The residues different from AAV3B of shuffled variants were 3D false-color mapped onto the crystal structure of AAV6 VP3. Light gray residues correspond to AAV3B amino acids while colored residues indicate surface exposed amino acids derived from other serotypes. With the exception of AAV3B color coding is as in (A) and (B).

### Evaluation of transduction efficiency of the new AAV variants on a panel of diverse cells lines

In order to evaluate if the AAV variants selected in human islets could also transduce other cell types, several primary cells as well as cell lines derived from human as well as animal sources were transduced with firefly luciferase rAAV vectors packaged with the two parental (AAV-DJ and AAV-LK03) as well as the three evolved AAV capsids (KP1, KP2, KP3) and analyzed for transduction efficiency using a luciferase assay (Figure 6A). As had been shown previously (Lisowski et al, 2014), AAV-LK03 did not transduce murine cells efficiently while AAV-DJ transduced all cell lines with high efficiency. Remarkably, the novel variants showed similar or slightly higher levels of transduction as compared to AAV-DJ on all cell types tested.

**Figure 6:**
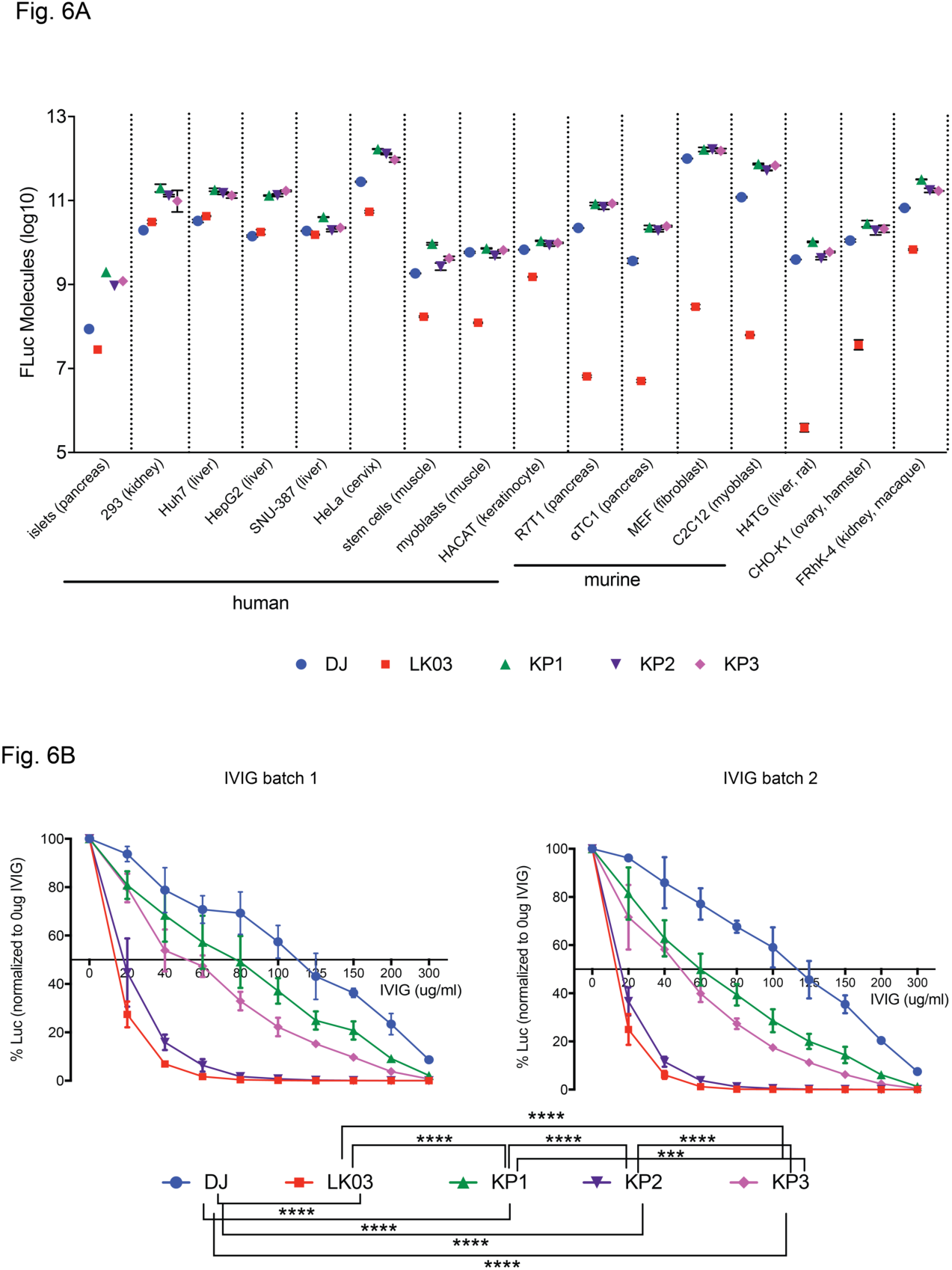
*In vitro* transduction experiments using rAAV-Firefly Luciferase Vectors. **(A)** Transduction efficiency of the novel capsids as well as AAV-DJ and AAV-LK03 capsids on a variety of human and non-human derived cell types. Cells were transduced with the different capsid containing rAAVs packaging a firefly luciferase expression cassette at a MOI of 1,000 in triplicates each (with the exception of islet cells) and cell lysates were analyzed 48 hrs post transduction in a luciferase activity assay. Two-fold dilutions of recombinant firefly luciferase enzyme were used to prepare a standard curve and raw luminescence units were calculated into luciferase molecules based on the standard curve. **(B)** Neutralization assay of rAAVs packaged with different capsids using dilutions of two different batches of pooled human immunoglobulin (IVIG). Huh-7 cells were transduced at a MOI of 100 with Firefly luciferase expressing rAAVs that had been pre-incubated with different concentrations of IVIG for 1 hr at 37°C. Luciferase activity in cell lysates was measured 24-hrs post transduction. Mean values of five replicates (obtained in two independent experiments) with standard deviations are shown for each sample. Only statistically significant differences are indicated in the legend below the graph. ***: p<0.001, ****: p<0.0001.

### Neutralization profile of the novel capsid variants

One of the limitations of many AAV vectors is the fact that the majority of patients exhibit pre-existing AAV neutralizing antibodies. Aiming to study the neutralization profile of our new variants, the novel capsid variants as well as AAV-DJ and AAV-LK03 were analyzed for sensitivity towards neutralization by two different batches of pooled human immunoglobulin (Figure 6B). The experiment was performed in seven biological replicates for each sample, split between two independent experiments. While AAV-DJ showed the highest resistance towards neutralization with ∼120 ug/ml IVIG needed for 50% inhibition, AAV-LK03 was more sensitive with 50% neutralization achieved with only ∼15 ug/ml. AAV-KP2 had a neutralization profile only slightly better than that of AAV-LK03 while AAV-KP1 and AAV-KP3 needed more IVIG to be neutralized (∼70 ug/ml and ∼50 ug/ml for AAV-KP1 and AAV-KP3, repectively). In summary, when pooled human IVIGs were used for neutralization, two of the novel variant capsids performed better than AAV-LK03, but less favourable than AAV-DJ.

### In vivo biodistribution of the AAV-KP variants in mice

Preclinical rAAV studies are routinely performed in mice, thus it is desirable that novel AAV capsids can be also used in this model prior to studies in human patients. As the novel capsid variants developed here had shown promise regarding transduction of a broad panel of different cell lines *in vitro,* we were interested to evaluate their transduction profile *in vivo* in mice. Mice were injected intravenously with firefly luciferase rAAV packaged with AAV8, AAV-DJ as well as AAV-KP1, KP2, and KP3 capsids and transgene expression was monitored over several weeks by live imaging (Figure 7, Figure S10A). AAV8 was included since AAV-LK03 does not transduce murine cells (Lisowski et al, 2014). As described previously (Palaschak et al, 2019), the majority of the injected AAVs was found to target the liver when delivering the virus intravenously. AAV-KP1 appeared to transduce mouse liver more rapidly than AAV-DJ, but expression levels were only slightly higher once steady state expression was achieved at later time points. Thirty-five days post injection, several organs were harvested from each mouse and analyzed for luciferase expression. With the exception of liver, low levels of expression with high intra-group variation were detected (Figure S10B). AAV-KP2 as well as AAV-KP3 injected mice had higher luciferase expression in heart tissue than AAV-DJ or AAV8 injected mice and all three variants had transduced spleen with higher efficiency than AAV-DJ or AAV8. Kidney tissue from mice injected with the novel variant AAVs also had higher luciferase expression than that from AAV-DJ injected mice, but levels were not significantly higher when compared to AAV8 injected mice. Vector genomes were quantified in the organs using qPCR. However, we were able to detect vector genome copies clearly above background only for the liver samples (Fig. S10C). Relative vector copy numbers correlated with the *in vivo* luciferase expression data.

**Figure 7:**
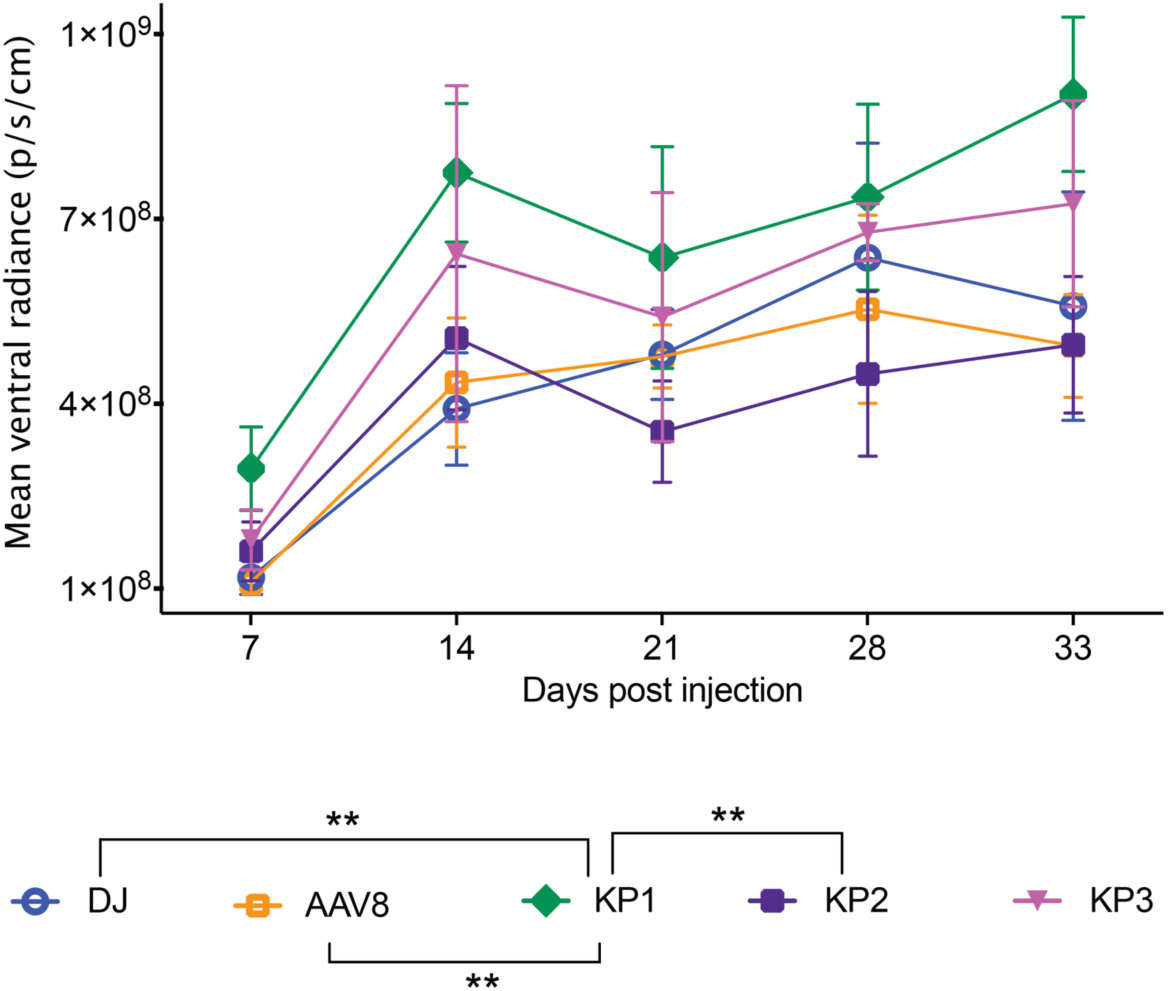
*In vivo* transduction efficiency of rAAVs packaged with the novel AAV8 and AAV-DJ capsids. Balb/C SCID mice were injected via tail vein with 2E10 vg each firefly luciferase expressing rAAV and luciferase expression in the livers was monitored over several weeks using live imaging after i.p injection of luciferin substrate. Four animals were injected for each group with the exception of the AAV8 group which contained 3 animals. The mean of each group’s mean ventral radiance is shown for each timepoint with standard deviations indicated. One animal from the AAV8 group was omitted from analysis due to a failed substrate injection. Only statistically significant differences are indicated in the legend below the graph. **: p<0.01

### Assessing functional human hepatocyte transduction in xenograft liver models in vivo

We transduced humanized FRG xenograft mice to assess the functional human hepatic transduction capabilities of KP1, the most promising of our novel capsids, in an appropriate *in vivo* setting. Mice were highly re-populated with human hepatocytes as shown by expression of high levels of human albumin (between 5.6 mg/ml and 8 mg/ml, data not shown). Humanized mice were administered Tomato Red expressing rAAV packaged with DJ, LK03, or KP1 capsids at a dose of 1E11vg/mouse via the intravenous injection, and assessed for expression of Tomato Red protein in human and mouse hepatocytes 14 days post-AAV administration (Figure 8). We observed that AAV-LK03 had high transduction efficiency in human hepatocytes, but not in mouse hepatocytes, while transduction with AAV-DJ was not specific towards human cells. Those findings are not unexpected as the LK03 capsid had been selected to be specific for transduction of primary human hepatocytes (Lisowski et al, 2014) while the DJ capsid was derived from an *in vitro* screen on human hepatoma cells in presence of neutralizing antibodies (Grimm et al, 2008), but has been reported to also transduce murine cells with high efficiency. AAV-KP1 was also found to transduce both human and mouse hepatocytes, however levels were higher than those found with AAV-DJ. In three of the four AAV-KP1 injected mice, transduction efficiency for human hepatocytes was similar to that found in AAV-LK03 injected mice. However, total transduction was higher in AAV-KP1 injected mice since unlike AAV-LK03, AAV-KP1 transduced both mouse and human cells.

**Figure 8:**
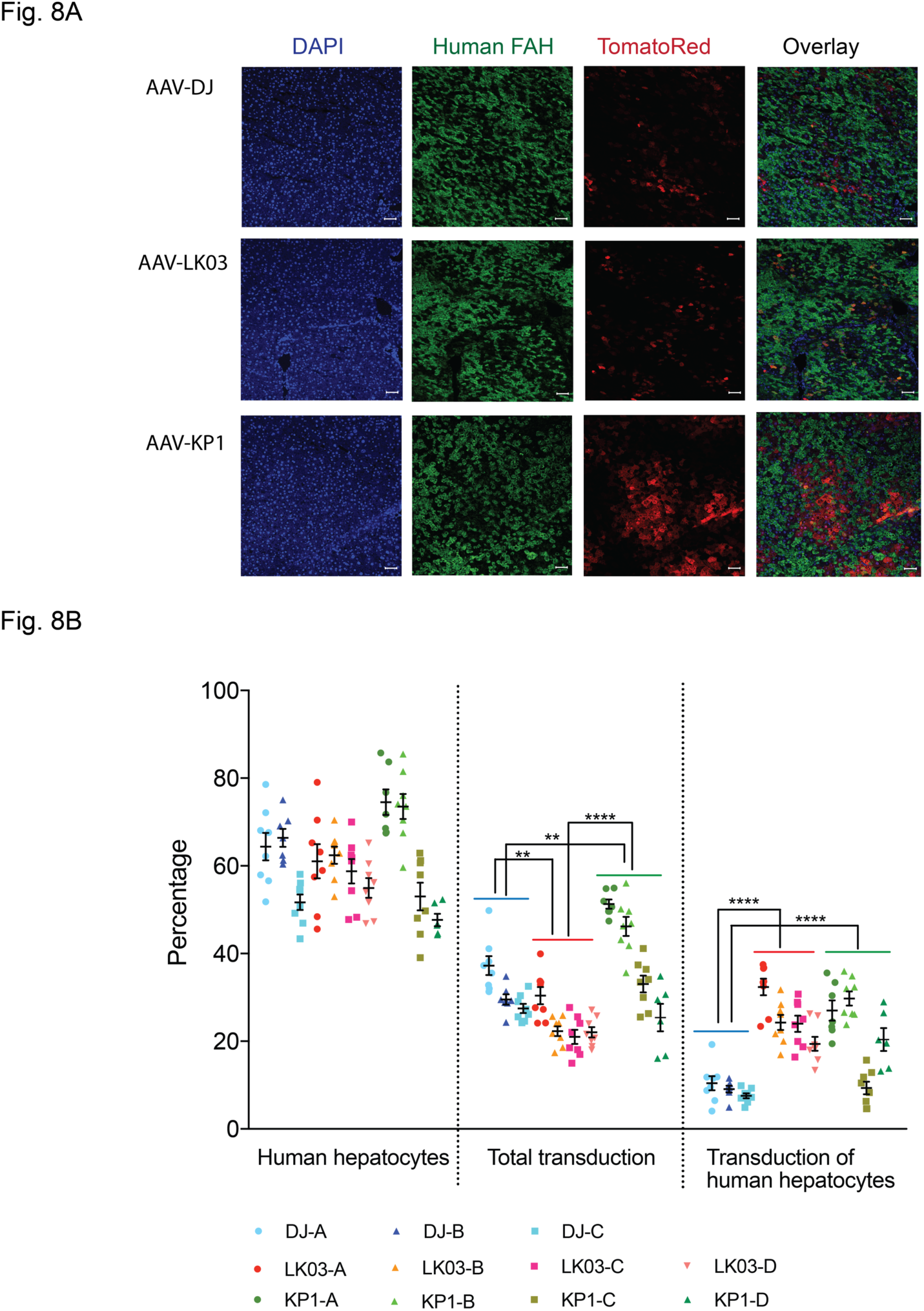
Validation and Quantification of Human Hepatocyte Transduction in Mice with Humanized Liver *in vivo*. **(A)** Representative immunofluorescence images from livers of mice injected with ssAAV-Td Tomato Red at 1E11 vg i.v. with varying capsid serotypes (3 mice for DJ, 4 mice each for LK03 and KP1). DAPI (blue), human-specific FAH (green), and Tomato Red (red) on liver sections. Scale bar, 50 µM. **(B)** Quantification of human hepatocyte repopulation levels, transduction efficiency for all hepatocytes as well as human hepatocytes. Each data point represents an area of interest for each mouse. A total of six to nine areas of interest for each mouse were scanned and analyzed. The mean and standard deviation for each mouse is indicated. Only statistically significant differences between the groups are shown in the graphs. **: p<0.01, ****: p<0.0001

## DISCUSSION

In this work we show for the first time that screening of highly diverse shuffled AAV capsid libraries that are tagged with unique short barcodes yielded highly functional AAV capsids for gene transfer into human islets as well as other clinically relevant cell types. The technology of tagging AAV capsids with individual barcodes had previously been described for *in vivo* tracking of variants derived from small AAV libraries or parental pools and has been termed AAV Barcode Seq (Adachi et al, 2014). Importantly, here we described the method for generation of barcoded capsid shuffled AAV libraries in great detail so that other researchers can easily apply this protocol in their own laboratory, either with genes from AAV or any other virus to generate libraries that can be screened for the desired properties. In addition to the protocol for library generation, we describe the production of a pool containing parental AAVs that proved to be useful as a parental control during the passaging regimen. This pool can be used in any library selection to optimzize the selection parameters to isolate AAV capsids with a desired property. The cost and time of the screening process can be minimized by predeterming the appropriate MOI and number of selection rounds for a particular screen. For previous library screening studies in our lab that did not employ barcoded libraries, four or more rounds of selection were routinely performed (Grimm et al, 2008; Lisowski et al, 2014; Paulk et al, 2018a; Paulk et al, 2018b). However, in the current study, we found that three rounds of selection were sufficient to enrich and amplify variants with improved transduction of the target cell type to a level where they could be isolated reliably. In addition, combining short DNA barcodes and single molecule DNA sequencing approaches will enable us to optimize other important parameters, such as the best MOI to use during the selection process, and the number of screening rounds necessary for efficient enrichment. When generating the AAV libraries from the plasmid libraries by transfection, we were careful not to overload the cells with library plasmid DNA by reducing the amount of AAV plasmid DNA almost 20-fold as compared to our routine protocol for virus production. It had been reported that a transfected copy number of 5,000 AAV genomes per cell is sufficiently low to minimize cross-packaging while still achieving high-titer libraries (Nonnenmacher et al, 2015). While the possibility of cross-packaging, which is characterized by sequence differences between the packaging capsid and the corresponding viral genome, cannot be excluded, several of the enriched capsids from our screen were found to exhibit the desired phenotype, suggesting that cross-packaging or mosaicism was not a major problem in our libraries. Moreover, when we had previously generated AAV libraries using high concentrations of AAV plasmid during transfection, we had been successful in generating variants with greatly improved phenotypes (Grimm et al, 2008; Lisowski et al, 2014; Paulk et al, 2018a; Paulk et al, 2018b), suggesting that for reasons unknown so far there seems to be a strong capsid genotype linkage in case of AAV packaging (Grimm & Zolotukhin, 2015). A recent report by the Grimm group laid to rest the concern that functionality of AAV capsid libraries generated for directed evolution studies might be severely compromised by inactivation of the assembly activating protein (AAP) in a large proportion of the chimeric variant pool (Herrmann et al, 2018b). Although this small protein had previously been described to play an important role in virus assembly (Grosse et al, 2017; Naumer et al, 2012; Sonntag et al, 2011; Sonntag et al, 2010), it appears that AAV is strikingly tolerant towards recombination within the AAP coding sequence. We had previously tested random capsids from other shuffled libraries for packaging efficiency and found that while most chimeric capsids produced rAAV with reduced titers as compared to AAV-DJ, very few capsids had a complete packaging defect. Indeed, it has been described that natural AAV serotypes 4, 5, and 11 do not require AAP for successful capsid assembly(Earley et al, 2017). The AAP sequences of all the chimeras selected in the present studies are chimeric and the rAAV titres obtained with two of the capsids were lower than those obtained with LK03. We did not evaluate if higher titer rAAV preparations can be obtained by supplementing wildtype AAP *in trans* as has been reported previously (Zinn et al, 2015) but this is certainly a possiblilty that can be explored to boost production yields.

When we subjected the libraries to stringent selection pressure on human islets, distinct capsid sequences were enriched after a few rounds of selection. Several of those have improved transduction efficiency for human islets as compared to the best known capsids to date – LK03 and DJ. The best performing capsids identified in the current study were at least 10-times more efficient at transducing human islet cells, particularly β-cells. In addition, when one of the variants was tested for transduction efficiency of hESC derived β-cells we were able to achieve similar levels of improvement. In addition, we confirmed prior observations that capsid variants AAV-DJ and AAV-LK03 were capable of transducing primary human islets with equal or better efficiency than AAV2 or AAV3B. While it is not clear which amino acid residues in the novel capsids are responsible for the improved islet phenotype we noticed several features in the improved capsid sequences that may confer enhanced transduction of islet cells as well as other cell types. It is possible that several surface exposed tyrosine residues that were described to trigger degradation of the capsids (Zhong et al, 2008) can be mutated to achieve further enhancement of transduction efficiency for the novel capsids.

Although the novel vectors described in this study mainly transduced the liver when delivered systemically, they could possibly be safely delivered in close proximity to islets using endoscopic retrograde cholangio-pancreatography (ERCP), a procedure that is routinely used in patients to examine the pancreatic and bile ducts (Hibberts & Barnes, 2003). Several recent studies have reported retrograde pancreatic intraductal delivery for safe and effective administration of rAAV vectors into the pancreas of mice (Jimenez et al, 2011; Mallol et al, 2017; Quirin et al, 2018). The vectors generated in our study could be used for gene therapy of diabetes by delivering transcription factors or small hairpin RNAs (shRNA) into pancreatic islet cells. The overexpression or inhibition of several transcription factors, such as Pdx1, Ngn3, MafA, Pax4, and Arx was found to effectively convert pancreatic islet progenitors and committed islet α-cells into β-cells (Chakravarthy et al, 2017; Collombat et al, 2009; Furuyama et al, 2019; Matsuoka et al, 2017; Wang et al, 2018; Xiao et al, 2018; Zhang et al, 2016). Moreover, expression of other factors, such as Igf1 and Follistatin has also shown promise in preserving β-cell mass or protecting newly transplanted donor islets from apoptosis (Mallol et al, 2017; Zhao et al, 2015).

When developing a vector for gene therapy, it is important to keep the goal of a clinical application in mind. Therefore, it is desirable to screen AAV capsid libraries in cells of human origin rather than in murine or other animal models. This has been clearly evidenced by a highly neurotropic AAV variant (AAV-PHP.B) that had recently been discovered through an *in vivo* library screen in mice (Deverman et al, 2016) and unfortunately was recently found to exhibit this enhanced tropism in mice only (Hordeaux et al, 2018; Matsuzaki et al, 2018). However, for the purpose of pre-clinical testing, it is desirable to develop capsids that can transduce murine cells in addition to human cells as this can accelerate the translation of different gene therapy approaches to the clinic. It is interesting that KP1 – despite its similarity to AAV3B and AAV-LK03 – transduced mouse hepatocytes as well as human hepatocytes in the humanized liver mouse model. This humanized model was used previously to select for several highly human liver AAV transducing vectors (Lisowski et al, 2014; Paulk et al, 2018b), one of which has been shown to provide robust human Factor VIII expression in an early human clinical trial (Doshi & Arruda, 2018; George, 2017).

However, all of the vectors selected in the previous screen showed poor transduction of mouse tissues or cells *in vitro* and *in vivo*. In contrast, the KP1 variant showed similar transduction of human hepatocytes in the chimeric mouse model but, in contrast to AAV-LK03, also showed efficient transduction of mouse liver. This is intriguing because some researchers have suggested that the transduction of human hepatocytes in this mouse model was exaggerated because of the relatively low transduction of mouse cells (Wang et al, 2015). If the data obtained from the human hepatocytes that had been engrafted in the mice is influenced by the degree of mouse hepatocyte transduction, perhaps the KP1 variant will provide even more robust transduction when tested in human trials where the liver cell population will be more homogenous. At a minimum, because this vector transduces both mouse and human liver, a surrogate capsid for preclinical testing is not required. Ultimately, these capsids are good candidates for future study in human clinical liver based gene therapy trials. In fact, because islet transduction will require localized delivery into the pancreatic duct, the KP variants could potentially be used in both applications.

The use of barcoded libraries for molecular evolution studies was found to be highly beneficial as high-throughput analysis of the barcodes of enriched variants is much more cost-effective and thorough than cloning and sequencing of the entire capsids. The libraries generated for this study are currently being evaluated in screens on a number of other target cells and may result in the discovery of other AAV capsid variants that are useful for other clinical gene therapy applications. The pool of parental barcoded AAVs proved to be a convenient and effective tool for validatation of the library selection parameters for the current study.

## METHODS

### Generation of an AAV vector containing a library of unique barcode sequences

A wildtype AAV2 vector in which the capsid coding sequences had been replaced by a PacI and AscI containing linker fragment (kindly provided by D. Grimm and S. Grosse, University of Heidelberg, Germany) was used as the starting material for construction of the barcoded AAV library. Two unique restriction sites (AgeI and EagI) were introduced just downstream of the cap polyadenylation signal by mutating two nucleotides in the original sequence (A to T and T to C at positions 6 and 24 when counting the last nucleotide of the cap polyA signal as starting point). Barcodes consisting of two stretches of 12 random nucleotides separated by a 20 nt long spacer sequence were generated as described previously(Adachi et al, 2014). Briefly, an oligonucleotide with an AgeI restriction site sequence on the 5’- end followed by 12 random nucleotides and the 20nt long spacer sequence (CTA AAC CGG TNN NNN NNN NNN NAC GGA AAT ACG ATG TCG GGA) was annealed to an oligonucleotide containing an Eag I site on the 5’ end followed by 12 random nucleotides as well as the antisense spacer sequence (TTC TCG GCC GNN NNN NNN NNN NTC CCG ACA TCG TAT TTC CGT) and extended using Klenow Polymerase devoid of exonuclease activity (NEB). Fragments were purified using a Qiaquick PCR purification kit (Qiagen) and subsequently digested with AgeI and EagI and purified using a Qiaquick PCR purification column. Vector was digested with the same restriction enzymes, dephosphorylated, phenol-chloroform purified and Ethanol precipitated. The optimum vector to insert ratio was evaluated by setting up several ligation reactions initially and testing for possible multiple barcode inserts by performing colony PCR with primers rightF (CGC GCC ACT AGT AAT AAA C) and QSeqRev (TAG AGC AAC TAG AGT TCG). For the scale-up reaction barcodes were ligated with the vector at a molar vector to insert ratio of 1 to 2.5 in a total volume of 30 ul, de-salted using Strataclean resin according to instruction (Agilent Technologies) and electroporated in 2 ul aliquots into DH10B-MegaX cells (Thermo Fisher). Electroporated cells were pooled and used to inoculate 500 ml LB-Amp medium. An aliquot was plated to assess library size and diversity. After 16 hrs in a 37°C shaker bacteria were harvested and plasmid DNA was isolated using a Megaprep kit (Qiagen). ITR integrity was confirmed by digestion with XmaI as well as with AhdI. Several individual clones from the test plate were sequenced to assess barcode diversity.

### Generation of capsid shuffled barcoded AAV libraries

DNAse I mediated family shuffling was essentially performed as described previously (Herrmann et al, 2018a; Kienle et al, 2012; Pekrun et al, 2002). Capsid sequences from 16 AAV serotypes (AAV1, AAV2, AAV3B, AAV4, AAV5, AAV6, AAV8, AAV9hu14, AAV12, AAV rhesus10, AAV porcine1, AAV porcine2, AAV bovine, AAV mouse1, AAV avian, AAV goat1) as well as from shuffled variants AAV DJ and AAV LK03 that had been selected in previous screens (Grimm et al, 2008; Lisowski et al, 2014) were used as parental sequences for the shuffling reactions. The capsid sequences had been obtained from various sources (D. Grimm, University of Heidelberg, Germany, Kay lab, Vector Core, Stanford). All sequences contained a PacI site immediately 5’ and an AscI site 3’ of cap and had been cloned into pBluescript. Prior to shuffling capsid sequences were amplified individually using primers located in the flanking pBluescript sequences (outer F: AAT TAA CCC TCA CTA AAG G, outer R: GTA ATA CGA CTC ACT ATA GGG C). Phusion Hot Start Flex polymerase (NEB) was used for amplification and 25 PCR cycles were employed (30 sec 98°C, 25 cycles of 10 sec 98°C, 15 sec 56°C, 1 min 15-sec 72°C, followed by 10 min 72°C). PCR products were purified using the Qiaquick PCR purification kit (Qiagen), all 18 capsids (for the 18 parent library) or 10 capsids (for the 10 parent library, AAV1, AAV2, AAV3B, AAV6, AAV8, AAV9hu14, AAV12, AAV rhesus10, DJ, LK03) were pooled in equimolar ratio and fragmented at room temperature (RT) using DNAse I (Sigma). At different incubation time points aliquots were analyzed on an 1.5% agarose gel while the reaction was temporarily stopped by incubation in a dry ice / ethanol bath. Incubation time and DNAse concentration was adjusted until the majority of the fragments ranged from 100 bp to 500 bp. The entire reaction was then loaded on an 1.5% agarose gel and fragments in the desired size range were electroeluted from the gel, purified using two rounds of phenol-chloroform purification followed by one round of chloroform purification and ethanol precipitated. DNA fragments were then re-assembled in a primer-less PCR using Phusion Hot Start Flex polymerase and the following cycling conditions: 30 sec 98°C, 40 cycles of 10 sec 98°C, 30 sec 42°C, 45 sec 72°C, followed by 10 min 72°C.

Full-length capsid sequences were amplified from the assembly reactions using primers rescueF (GTC TGA GTG ACT AGC ATT CG) and rescueR (GTC TAC TGA AGC TCA CTG AG) and the following cycling conditions: 30 sec 98°C, 25 cycles of 10 sec 98°C, 15 sec 57°C, 1 min 15 sec 72°C, followed by 10 min 72°C. Amplicons were diluted 4-fold with fresh PCR mix and subjected to one additional cycle with a 10 min extension to fill up the ends. After concentrating PCR products using a PCR purification kit (Qiagen) they were digested with PacI and AscI and ligated into the BC library vector that had been treated with PacI, AscI, dephosphorylated and phenol-chloroform purified. Ligation reactions were de-salted using the Strataclean resin, electroporated into MegaX DH10B cells and expanded in liquid culture as described above for generation of the BC library vector. Small aliquots of transformed cells were plated to assess library size. Plasmid DNA was extracted from random colonies and *cap* and BC sequences were determined by Sanger sequencing using primers capF (TGG ATG ACT GCA TCT TTG AA), capF2 (ATT GGC ATT GCG ATT CC), and QSeqRev (TAG AGC AAC TAG AGT TCG).

AAV libraries were generated in HEK 293T cells using the calcium phosphate transfection method. Compared to the regular protocol the amount of transfected library plasmid DNA was reduced almost 20-fold to approximately 5000 copies per cell to minimize the likelihood of cross-packaging events taking place during AAV production. Briefly, 25 T225 flasks were seeded with 8E06 cells per flask in 40 ml media two days prior to transfection. On the day of transfection cells were between 80% and 90% confluent. 20 ml of media per flask was replaced with fresh media 1.5 hrs prior to transfection and a mixture of 40 ug pAd5 helper plasmid and 2 ug library plasmid in 4 ml 300mM CaCl_2_ per T225 was prepared. Equal amounts of CaCl_2_ / DNA mix and 2xHBS (280 mM NaCl, 50 mM HEPES pH 7.28, 1.5 mM Na_2_HPO_4_, pH 7.12) were mixed and 8 ml of the mixture was added to each flask. After 3 days cells were detached with 0.5 ml 500 mM EDTA each flask and the cell pellet was resuspended in Benzonase digestion buffer (2 mM MgCl_2_, 50 mM Tris-HCl, pH 8.5). AAVs were released from the cells by submitting them to three freeze-thaw cycles, non-encapsidated DNA was removed by digestion with Benzonase (200 U/ml, 1 hr 37°C), cell debris was pelleted by centrifugation, followed by another CaCl_2_ precipitation step (25 mM final concentration,1 hr on ice) of the supernatant and an AAV precipitation step using a final concentration of 8% PEG-8000 and 625 mM NaCl. Virus was resuspended in HEPES-EDTA buffer (50 mM HEPES pH 7.28, 150 mM NaCl, 25 mM EDTA) and mixed with CsCl to a final refractory index (RI) of 1.371 followed by centrifugation for 23 hrs at 45000 Rpm in a ultracentrifuge. Fractions were collected after piercing the bottom of the centrifuge tube with a 18 gauge needle and fractions ranging in RI from 1.3766 to 1.3711 were pooled and adjusted to an RI of 1.3710 with HEPES-EDTA resuspension buffer. A second CsCl gradient centrifugation step was carried out for at least 8 hrs at 65000 Rpm. Fractions were collected and fractions with an RI of 1.3766 to 1.3711 were dialyzed overnight against PBS, followed by another 4 hr dialysis against fresh PBS and a 2 hr dialysis against 5% sorbitol in PBS. All dialysis steps were carried out at 4°C. Virus was recovered from the dialysis cassette and pluronic F-68 was added to a final concentration of 0.001%. Virus was sterile-filtered, aliquoted, and stored in aliquots at -80°C. Genomic DNA was extracted from 10 ul of the purified virus using the MinElute Virus Spin Kit (Qiagen Cat#57704), and the viral genome titer was determined by qPCR using an AAV2 rep gene specific primer probe set (repF: TTC GAT CAA CTA CGC AGA CAG, repR: GTC CGT GAG TGA AGC AGA TAT T, rep probe: TCT GAT GCT GTT TCC CTG CAG ACA).

### Generation of barcoded parental AAV pools

Capsids from all 18 parental AAVs were cloned into the BC library vector using PacI and AscI restriction sites. Each parental AAV contained a unique barcode sequence as confirmed by sequence analysis and between 2 and 6 T225 with 293T cells were transfected with each parental AAV (37.5 ug AAV plasmid and 37.5 ug pAd5 helper plasmid per T225). Crude lysates of each barcoded parental cap AAV were generated and 2.8E12 vg of the 10 parents or 1.1E12 vg of the 18 parents were pooled to generate the 10 parent and the 18 parent mix respectively. AAV pools were purified by double CsCl gradient centrifugation as described above.

### Sequence contribution analysis of evolved AAV capsids

Contigs were assembled using Sequencher 5.3 software and aligned using the Muscle multiple sequence alignment software (MacVector, Version 14.5.3). Xover 3.0 DNA/protein shuffling pattern analysis software(Huang et al, 2016) was used to generate parental fragment crossover maps of shuffled variants. Each parental serotype was color coded as indicated in the figures.

### PacBio sequencing of AAV pools and libraries

For the 10 parent library as well as the 18 parent pool a 2.4 kb fragment containing the capsid as well as the BC sequences was amplified using capF and QSeqRev from extracted viral genomes, loaded onto a 1% agarose gel, visualized by staining with SybrSafe, and gel-purified using a gel extraction kit (Qiagen). The 10 parent library was also assessed at the plasmid level prior to generating the AAV library using restriction enzymes (PacI and XbaI) to release the capsid sequences and gel purified as described above for the amplified capsid sequences. Library preparation and Pacific Biosciences (PacBio) sequencing were performed at the University of Washington PacBio Sequencing Service. Briefly, SMRT bell libraries were prepared following the “Procedure and Checklist-2 kb Template Preparation and Sequencing” protocol from PacBio using the SMRTbell Template Prep Kit v1.0 (PacBio Cat#100-259-100). PacBio ‘Binding and Annealing’ calculator was used to determine appropriate concentrations for annealing and binding of SMRTbell libraries. SMRTbell libraries were annealed and bound to P6 DNA polymerase for sequencing using the DNA/Polymerase Binding Kit P6 v2.0 (PacBio Cat#100-372-700). Bound SMRTbell libraries were loaded onto SMRT cells using standard MagBead protocols and the MagBead Buffer Kit v2.0 (PacBio Cat#100-642-800). The standard MagBead sequencing protocol was followed with the DNA Sequencing Kit 4.0 v2 (PacBio Cat#100-612-400, also known as P6/C4 chemistry). Sequencing data was collected for 6 hr movie times with ‘Stage Start’ not enabled. Circular consensus sequence (CCS) reads were generated using the PacBio SMRT portal and the RS_ReadsOfInsert.1 protocol, with filtering set at Minimum Full Pass = 3 and Minimum Predicted Accuracy = 95%.

### Bioinformatic assessment of PacBio sequence reads

CCS reads with full capsid sequence lengths from 2,250-2,380 nucleotides were included in downstream bioinformatics analyses. Indels in CCS reads were corrected using an in-house algorithm that first assesses parental fragment identity using Xover 3.0 DNA/protein shuffling pattern analysis software. Once the parental identity of each crossover fragment was determined, this information was used to determine indels for correction. Single nucleotide polymorphisms (SNPs) that did not result in indels were maintained. The SNP error rate with the PacBio platform is 1.3-1.7%. SNP frequencies above this rate range were assumed to have arisen from *de novo* mutations. Corrected sequences in FASTA format were then aligned with MUSCLE. Phylogenetic analyses were conducted using the maximum-likelihood method in RAxML(Stamatakis, 2015; Stamatakis et al, 2005).

### False-colored structural capsid mapping

Chimeric capsids (VP3 sequences only) were false-color mapped onto the AAV6 structure 4V86 (Xie et al, 2011) using PyMOL Version 2.3.0. Only surface exposed amino acids that are different from the closest parental AAV3B are shown. With the exception of AAV3B which is shown in gray all mapped amino acid residue colors correspond to parental serotype colors used in the crossover and enrichment figures.

### Conservation and enrichment calculations

Amino acid conservation for each position was calculated using the alignment profile obtained with MacVector version 14.5.3. Average conservation values were calculated for stretches of 30 amino acid residues and were used to generate the graphs. Percent parental conservation was determined using an in-house algorithm that identifies the percentage of each parent on each aligned position in the shuffled library. The maximum square size indicates that 100% of variants share that amino acid from that parent at that position. All other square sizes are proportional to the percent of variants from 0-100% that have that amino acid at that position from that parent. Enrichment scores were calculated for each amino acid position in the sequence of each chimera by comparison of sequences from parental serotypes based on maximum likelihood. Xover version 3.0 (Huang et al, 2016) was used to generate a crossover data analysis set for each chimera. Excel version 16.20 was used to convert those data into enrichment scores. Library parents are depicted in different colors as shown.

### Statistics

Statistical analyses were conducted with Prism v7.0d. Experimental values were assessed via two-way ANOVA using Tukey’s multiple comparisons test. *P* values <0.05 were considered statistically significant.

### High throughput sequencing of AAV barcodes

Barcode sequences were amplified with indexed primers (F: AAT GAT ACG GCG ACC ACC GAG ATC TAC ACT CTT TCC CTA CAC GAC GCT CTT CCG ATC T **(I)** CGC GCC ACT AGT AAT AAA C and R: CAA GCA GAA GAC GGC ATA CGA GAT CGG TCT CGG CAT TCC TGC TGA ACC GCT CTT CCG ATC T **(I)** TAG AGC AAC TAG AGT TCG, with the indices (**I**) containing between 4 and 6 nucleotides), gel-purified from 2% SybrSafe containing agarose gels, pooled (up to 30 samples), and sequenced on a MiSeq instrument. The number of PCR cycles was minimized to avoid amplification bias and was dependent on the concentration of input AAV genomes as determined by rep qPCR. The following cycling conditions were used: 2 min 98°C, 15 to 30 cycles of 15 sec 98°C, 15 sec 50°C, 20 sec 72°C, with a final 15-min extension at 72°C. Phusion Hot Start Flex (NEB) was used for all amplifications.

### Cell culture conditions

#### Human islet cultures

Human pancreatic islets from deceased non-diabetic organ donors were provided by the Integrated Islet Distribution Program (IIDP) or the University of Alberta through the Stanford Islet Research Core and cultured in CMRL-1066 with 10% FBS, Pen-Strep, 1% Insulin Transferrin Selenium (Thermo Fisher), 1mM sodium pyruvate, 2mM Glutamax, 2.5mM HEPES. Ultra-low attachment dishes were used for all islet cell culture experiments.

*hESC derived β-cells:* Mel1 INS^GFP/W^ human embryonic stem cell (hESC)s were obtained from S. J. Micallef and E. G. Stanley (Monash Immunology and Stem Cell Laboratories, Australia). Cells were maintained and propagated on mouse embryonic fibroblasts (MEFs) in hESC media [DMEM/F12 (Gibco) with 10% KSR (Gibco), 10 ng/ml FGF-2 (R&D Systems)]. A stepwise differentiation of hESC toward β cells was carried out following the protocol described previously(Nair et al, 2019). Briefly, confluent hESC were dissociated into single-cell suspensions using TrypLE (Gibco), counted and seeded at 5.5 × 10^6^ cells per well in 6-well suspension plates in 5.5 ml hESC media supplemented with 10 ng /ml activin A (R&D Systems) and 10 ng/ ml heregulinB (Peprotech). The plates were incubated at 37 °C and 5% CO_2_ on an orbital shaker at 100 rpm to induce 3D sphere formation. After 24 hours, the spheres were washed with PBS and resuspended in day 1 media. From day 1 to day 20, media was changed every day. Media compositions are as follows: Day 1: RPMI (Gibco) containing 0.2% FBS, 1:5,000 ITS (Gibco), 100 ng /ml activin A and 50 ng /ml WNT3a (R&D Systems). Day 2: RPMI containing 0.2% FBS, 1:2,000 ITS and 100 ng /ml activin A. Day 3: RPMI containing 0.2% FBS, 1:1,000 ITS, 2.5 μM TGFbi IV (CalBioChem) and 25 ng /ml KGF (R&D Systems). Day 4–5: RPMI containing 0.4% FBS, 1:1,000 ITS and 25 ng/ ml KGF. Day 6–7: DMEM (Gibco) with 25 mM glucose containing 1:100 B27 (Gibco) and 3 nM TTNPB (Sigma). Day 8: DMEM with 25 mM glucose containing 1:100 B27, 3 nM TTNPB and 50 ng /ml EGF (R&D Systems). Day 9–11: DMEM with 25 mM glucose containing 1:100 B27, 50 ng /ml EGF and 50 ng/ ml KGF. Day 12–20: DMEM with 25 mM glucose containing 1:100 B27, 1:100 Glutamax (Gibco), 1:100 NEAA (Gibco), 10 μm ALKi II (Axxora), 500 nM LDN-193189 (Stemgent), 1 μm Xxi (Millipore), 1 μM T3 (Sigma-Aldrich), 0.5 mM vitamin C, 1 mM N-acetyl cysteine (Sigma-Aldrich),10 μM zinc sulfate (Sigma-Aldrich) and 10 μg /ml of heparin sulfate. At day 20, the spheres were collected, incubated with Accumax (innovative cell technologies) for 10 min at 37 °C and dissociated into single cells. Live GFP-high cells were sorted on Aria II at low flow rates and reaggregated in Aggrewell-400 (StemCell Technologies) at 1,000 cells per cluster in CMRL containing 10% FBS, 1:100 Glutamax (Gibco), 1:100 NEAA (Gibco), 10 μm ALKi II (Axxora), 0.5 mM vitamin C, 1 μM T3 (Sigma-Aldrich), 1 mM N-acetyl Cysteine (Sigma-Aldrich), 10 μM zinc sulfate (Sigma-Aldrich) and 10 μg /ml of heparin sulfate. At day 23, the reaggregated enriched β-clusters (eBCs) were transferred from Aggrewells and placed on orbital shakers at 100 rpm, and further cultured for 6 days. Media was changed every third day following reaggregation.

#### Human skeletal muscle stem cell and myotube cultures

A pool of primary muscle stem cells isolated from 6 individual donors (kind gift from G. Charville, Stanford) was frozen at an early passage and aliquots were used for experiments. Plates were coated with extracellular matrix protein (Sigma) at 1:500 in DMEM with 1% penicillin/streptomycin. The hMuSC medium was a 1:1 mixture of DMEM:MCDB media supplemented with 20% FBS, 1% insulin-transferrin-selenium, 1% antibiotic/antimycotic, and 10 μM p38i (Cell Signaling Technology Cat#SB203580) to maintain the stem state as described (Charville et al, 2015). Media for differentiating primary hMuSCs into myotubes lacked p38i and included a 2% horse serum starve instead of 20% FBS for 7 days. All media was changed every two days.

#### Mouse skeletal muscle myoblast cultures

Wild-type C2C12 mouse myoblasts (ATCC Cat# CRL-1772) were maintained in DMEM supplemented with 10% FBS and 1% antibiotic/antimycotic. *293 and 293T cell line cultures.* HEK 293 cells (ATCC Cat# CRL-1573) and HEK 293T cells (ATCC Cat#CRL-3216) were cultured in DMEM with 10% FBS, 2 mM glutamine, 1% antimycotic-antibiotic, 11 mM HEPES pH 7.28 and 1 mM sodium pyruvate.

#### HeLa cell cultures

HeLa cells (ATCC Cat# CCL-2) were cultured in DMEM with 10% FBS, 2mM glutamine, 1% antimycotic-antibiotic.

#### Mouse pancreatic β-cell cultures

R7T1 cells (kind gift from H. Moeller, Stanford) were cultured in DMEM with 10% FBS, 2mM glutamine, 1% antimycotic-antibiotic, 1 ug/ml Doxycyclin.

#### Mouse pancreatic α-cell cultures

Alpha TC1 clone 6 cells (ATCC Cat# CRL-2934) were cultured in DMEM with 10% FBS, 2mM glutamine, 1% antimycotic-antibiotic, 15 mM HEPES, 0.1 mM NEAA.

#### Rat hepatoma cell cultures

H4TG cells (ATCC Cat# CRL-1578) were cultured in DMEM with 10% FBS, 4 mM glutamine, 1% antimycotic-antibiotic.

#### Human hepatocellular carcinoma cell cultures

SNU-387 cells (ATCC Cat# CRL-2237) and HepG2 (ATCC Cat# HB-8065) were cultured in RPMI with 10% FBS, 2mM glutamine, 1% antimycotic-antibiotic, 1% non-essential amino acids. HuH7 cells were cultured in DMEM with 10% FBS, 2 mM glutamine, 1% antimycotic-antibiotic, 1% non-essential amino acids.

#### Human keratinocyte cell cultures

HaCaT cells (Boukamp et al, 1988) (kind gift from A. Oro, Stanford) were cultured in DMEM with 10% FBS, 2 mM glutamine, 1% penicillin/streptomycin. *Hamster ovary cell cultures*. CHO-K1 cells (ATCC Cat# CCL-61) were cultured in Ham’s F12 with 10% FBS, 1% penicillin/streptomycin.

#### Rhesus macaque kidney cell cultures

FRhK-4 cells (ATCC Cat# CRL-1688) were cultured in DMEM with 10% FBS, 2mM glutamine, 1% penicillin/streptomycin.

#### Mouse fibroblast cell cultures

Primary mouse embryonic fibroblasts derived at E14 were cultured in DMEM with 10% FBS, 2 mM glutamine, 1% non-essential amino acids, 1% antimycotic-antibiotic, 55 uM β-Mercaptoethanol.

### Selection of AAV libraries on human islets

Islets were left in a 10 cm Petri Dish with 10 ml complete media to recover overnight prior to AAV infection. Islets were infected either intact or were dissociated into single cell suspensions using Accumax prior to infection (1 ml Accumax per 1000 islet equivalents [IEQ]). Approximately 300 IEQ or 1.7E05 dispersed islet cells were seeded in several wells of an ultra-low attachment 24-well plate, infected with various MOIs of either the 18 parent AAV mix or the AAV libraries and incubated in a 37°C incubator for 6 hrs. After two PBS washes to remove left over input virus cells were super-infected with human adenovirus 5 obtained from ATCC (Cat# VR-5). For intact islets 8E07 PFU were used, for dissociated islet cells 4E07 PFU were added into 1 ml media per well. After 4 days at 37°C the cells and supernatant were harvested, subjected to 3 freeze-thaw cycles and incubated for 30 min at 65°C to inactivate Ad5. Cell debris was removed by centrifugation (2 min, 10,000xg) and viral genomes were isolated from 100-ul clarified supernatant for titration by qPCR using a rep primer-probe set. For subsequent rounds of passaging similar MOIs as for the initial infections were used if sufficiently high titers were achieved. When titers were low a maximum volume of 200 ul was used for infection.

### Vector plasmids

A self-complementary rAAV vector expressing GFP under control of a CAG promoter (pscAAV-CAG-GFP, Addgene, Cat#83279) was generated by replacing the CMV promoter in plasmid pscAAV-GFP (gift from John T Gray, Addgene, Cat#32396) with the CAG promoter derived from pAAV-CAG-GFP (gift from Edward Boyden, Addgene, Cat#37825). A single stranded rAAV vector expressing Firefly luciferase (FLuc) under control of the CAG promoter (pAAV-CAG-FLuc, Addgene, Cat#83281) was generated by replacing the GFP sequences in plasmid pAAV-CAG-GFP with Firefly luciferase sequences obtained from plasmid pAAV-EF1α-FLuc-WPRE-HGHpA (Addgene, Cat#87951). A single stranded rAAV vector expressing codon diversified Tomato Red was a gift from Edward Boyden (pAAV-CAG-tdTomato, Addgene, Cat#59462).

### Recovery and evaluation of enriched AAV capsid sequences

Capsid sequences were amplified from viral genomes after the third round of selection using primer capF and a reverse primer containing the respective BC specific sequence on its 3’ end. The right BC was chosen to be included in the primer sequences so that the left BC served as a control of specific amplification of the desired variant. The number of PCR cycles was adjusted according to viral titer and frequency of the specific variant in the viral pool. The following amplification parameters were used: 2 min 98°C, 25 to 30 cycles of 15 sec 98°C, 20 sec 61°C, 2 min 72°C, with a final 10 min extension at 72°C. PCR products were gel purified, TOPO cloned and sequenced. At least three clones for each BC were sequenced to ensure that the left BC sequence matched the sequence obtained by BC NGS and to ensure that the sequences were identical. For most of the capsids that were amplified using the BC specific reverse primer we found clones that had more differences from the consensus sequence than it would be expected when using a proofreading polymerase. Since most of the sequence differences were found in the 5’ end and both BC sequences were correct we believe this observation was likely due to template switching after incomplete extension (Chakravarti & Mailander, 2008; Odelberg et al, 1995; Paabo et al, 1990) and may be alleviated by optimizing PCR conditions and enzymes. In the cases where this was observed a larger number of clones (up to 10 clones) was analyzed and the most abundant capsid sequences were used for vectorization. The capsids were used to package a self-complementary CAG promoter driven GFP expression vector by calcium phosphate triple transfection. For each T225 flask 25 ug sc CAG-GFP transfer vector, 25 ug packaging plasmid, and 25 ug pAd5 helper plasmid was used. Crude cell lysates were generated, rAAV titers determined by qPCR using a GFP specific primer-probe set, and tested for transduction efficiency of dissociated human islet cells using an MOI of 1K. 48-hrs post transduction the re-aggregated pseudo islets were dissociated into single-cell suspensions by incubation with Accumax followed by treatment with Dispase. The number of GFP expressing cells was evaluated using a BD FACS Calibur instrument and FlowJo software Version 10 was used to analyze and graph data. Selected capsid variants were used to generate CsCl gradient purified vector preparations packaging different expression vectors.

### Evaluation of cell type specific transduction efficiency of capsid variants

The cold transduction method was performed for those studies. Briefly, approximately 300 intact islets were resuspended in 100 ul CMRL with 2% FBS, rAAV was added at an MOI of 10K (assuming 1,000 cells per islet), and the mixture was incubated on ice while gently rocking on a horizontal shaker in the cold room. After 2 hrs 1 ml pre-warmed complete media (CMRL-1066 with 10 mM HEPES, 0.5% human serum albumin, 2% FBS, 10 mM nicotinamide, 1% antimycotic-antibiotic, 1% Glutamax) was added to each sample and islets were incubated on 24-well ultra-low attachment plates. Media was replaced after 2 days and islets were harvested, dissociated, and analyzed by FACS as described previously using surface antibodies to subdivide into α-, β-, and non-α-/non-β-cells(Dorrell et al, 2008).

### Evaluation of rAAVs for transduction efficiency on hESC derived β-cells

Recombinant AAVs were mixed with 800,000 GFP-high cells sored from 20 spheres at an MOI of 10, 100, 1000 and reaggregated in Aggrewell-400 in CMRL containing 10% FBS, 1:100 Glutamax, 1:100 NEAA, 10 μm ALKi II, 0.5 mM vitamin C, 1 μM T3, 1 mM N-acetyl Cysteine, 10 μM zinc sulfate and 10 μg /ml of heparin sulfate. Media was replaced after 3 days when the reaggregated eBCs were transferred into 6 well suspension plates. They were placed on orbital shakers at 100 rpm, and further cultured for 3 days. Subsequently, the eBCs were dissociated, fixed, permiabilized and stained for anti-human C-peptide abtibody (1:200), and anti-human RFP antibody (Rockland, 1:500) for analysis on LSRFortessa X20 Dual, as described previously(Russ et al, 2015). Data were analyzed with Flowjo software. Anti-human C-peptide antibody was conjugated in-house using the Molecular Probes Antibody Labeling Kits according to manufacturer’s instructions. Live images were taken using Leica DMI4000 B.

### Evaluation of the variants for transduction efficiency on a variety of cell lines

Capsid sequences of AAV DJ, AAV LK03, as well as the variants AAV KP1, KP2, and KP3 were used to package a single stranded CAG-Firefly Luciferase vector. Recombinant AAV preparations were double CsCl purified and used to transduce a variety of human and mouse primary cells and cell lines at an MOI of 1,000 in triplicates. Except for the differentiated human muscle cells all cells were seeded one day prior to transduction on 48-well plates so that they were about 60-70% confluent at the time of transduction (seeding density of 20,000-80,000 per well, depending on size and proliferation rate). Cells were lysed and assayed for luciferase activity using the Luciferase Assay Kit (Promega) 48 hrs post transduction. Purified recombinant luciferase protein (Promega) was used to generate a standard curve.

### Neutralization assay

Two different batches of pooled human immunoglobulin fractions (IVIG, Baxter) were used to evaluate the novel variants for sensitivity to neutralizing antibodies. Neutralization assays were essentially performed as described(Meliani et al, 2015). Briefly, IVIG preparations were diluted in complement inactivated FBS and incubated for 1 hr at 37°C with 2E08 vector genomes of each ssCAG-FLuc vector packaged with the different capsids in a total volume of 100 ul. Huh7 cells that had been seeded on 48-well plates the day before (5E04 per well) were transduced with the virus-IVIG mixtures in triplicates (22.5 ul each well, corresponding to MOI of ca 100) and luciferase activity in the cell lysates was determined 24 hrs later.

### Mice

*Fah/Rag*2*/Il2rgc* (FRG) deficient female mice on a NOD-strain background (FRG/N) were housed and maintained in specific-pathogen-free barrier facilities at Oregon Health & Science University. FRG/N mice were maintained on irradiated high-fat low-protein mouse chow (Lab Diet Cat#Picolab-5LJ5) *ad libitum* to decrease flux through the tyrosine pathway. Beginning on the day of transplantation, FRG/N mice were maintained for 1 week on acidified water to prevent bacterial growth. The following week, mice were switched to 1 week of 8 mg/L SMX-TMP antibiotic water (supplemented with 0.7 mol/L dextrose for palatability). Thereafter, FRG/N mice were cycled on and off 1 mg/L NTBC water as described. Female Balb/C SCID mice between 6 and 8 weeks of age were purchased from The Jackson Laboratories (Cat#001803) for imaging studies. The Institutional Animal Care & Use Committees of Stanford University and Oregon Health & Science University approved all mouse procedures.

### Hepatocyte transplantation

Donor human hepatocytes for transduction studies were acquired from BioreclarnationIVT (Lot#QIE). Weanling FRG/N mice were pre-conditioned with administration of recombinant human adenovirus expressing urokinase (5E10 PFU retroorbitally) 24 hrs prior to transplant to promote human cell engraftment. Between 5E05 and 1E06 human hepatocytes were injected intrasplenically into anesthetized recipient FRG/N mice and cycled on/off NTBC to promote human hepatocyte engraftment and expansion. Broad-spectrum antibiotic (Ceftiofur 4 mg/kg) was given by intraperitoneal injection immediately prior to surgery and for two days following surgery. Six months post-transplant, circulating human albumin levels as measure of engraftment were determined with the Bethyl Quantitative Human Albumin ELISA kit (Cat#E88-129).

### *In vivo* transduction experiments

For evaluation of wildtype mouse liver transduction efficiency white Balb/C SCID mice were injected with 2E10 vector genomes of each CAG-Firefly Luciferase vector via normodynamic intravenous lateral tail vein injections. AAV8 was used in place of LK03 as this capsid had previously been shown to be highly human specific. Mice were monitored for Luciferase activity in the liver once a week by intraperitoneal injection of 150 ug per g body weight D-Luciferin (Biosynth Cat#L-8220) and ventral luciferase readings using an Ami Imaging System. On day 35 mice were sacrificed and various organs were recovered (liver, pancreas, heart, lung, spleen, brain, and kidney). Organs were homogenized in Passive lysis buffer (Promega) using a Bullet Storm Homogenizer and luciferase activity from 1 mg each tissue sample was measured as described above. Genomic DNA was isolated from 10 mg each tissue sample and vector copy numbers were determined using qPCR. Primers for Luciferase were FLuc F: CAC ATA TCG AGG TGG ACA TTA C and FLuc R: TG TTT GTA TTC AGC CCA TAG. Mouse actin primers (m-actF: CCT GTA TGC CTC TGG TCG TA and m-actR: CCT CGT AGA TGG GCA CAG T) were used for normalization.

For evaluation of human hepatocyte transduction *in vivo* humanized FRG/N mice (3 mice for DJ, 4 mice each for LK03 and KP1) were injected intravenously with 1E11 ssCAG-Td Tomato Red vector genomes pseudotyped with DJ, LK03, or KP1 capsids and maintained on 1 mg/L NTBC during this 14 day transduction. Livers were harvested under inhalation isoflurane anesthesia. Liver tissue was cut into several 2×5-mm pieces from several lobes and fixed in 10x volume of 4% PFA for 5 hrs at 25°C protected from light. Fixed tissue was washed 1X in PBS and put through a sucrose cryoprotection and rehydration series (10% w/v sucrose for 2 hrs at 25°C, 20% w/v sucrose overnight at 4°C, 30% w/v sucrose for 4 hrs at 25°C). Liver pieces were rinsed in PBS, blotted dry and mounted in cryomolds (Tissue-Tek Cat#4557) with OCT (Tissue-Tek Cat#4583) and frozen in a liquid nitrogen-cooled isopentane bath. Labeled cryomolds were wrapped in aluminum foil and placed at -80°C until sectioning.

### Liver immunohistochemistry

Each liver sample with four to five lobes was cut in a microtome at 5 µM per section. Slides were fixed in methanol for 1 min at -20°C and air dried at RT. All following steps were done at RT except for notification. Slides were washed in PBT (PBS + 0.1% Tween20) for 3×5 min, permeabilized in 0.3% Triton-X-100 in PBS for 1×10 min, and washed in PBT for 2×5 min. Blocking was performed in PBT + 10% normal donkey serum (Cat. no. ab7475; Abcam) for 1 hr in a humidified chamber. A rabbit anti-human FAH antibody (Cat. no. HPA041370; Sigma) was added in PBT at 1:200 and incubated overnight at 4°C. Post-staining wash was conducted in PBT for 3×5 min. A secondary antibody for donkey anti-rabbit Alexa Fluor 488 IgG antibody (Cat. no. A-21206; ThermoFisher) was added in PBT (1:500, with DAPI at 80 ng/mL) and incubated at RT for 1 hr. Slides were washed in PBT for 3×5 min and PBS for 1×5 min, followed by mounting with ProLong Gold Antifade Reagent (Cat. no. 9071S; Cell Signaling). Antibody validity controls included secondary-only staining and demonstration on positive control human liver tissue sections (Cat. no. HF-314; Zyagen) and negative control untreated mouse liver sections (Cat. no. MF-314-C57; Zyagen). Imaging was performed on an inverted Zeiss laser scanning confocal microscope (LSM 880) by a 20× objective with the Zen Pro software. AAV-RFP signals were scanned and captured directly. Quantification of human hepatocyte transduction was done using the Volocity software (v6.3) and confirmed with counts by eye. Briefly, six to nine different areas of interests across different lobes from different sections of each liver sample were scanned, counting ∼1,000 cells on average per section. Areas with roughly 30%-80% of FAH-positive staining were chosen for analysis. The percentage of human hepatocytes per scanned area was presented as the number of cells with Alexa Fluor 488 signals (FAH-positive) divided by the number of cells with DAPI signals (total number of cells). Total transduction efficiency was calculated by dividing the number of cells with RFP signals (AAV-positive) by the number of cells with DAPI signals. The overlap of these two numbers represented the transduction efficiency for human hepatocytes for different AAV serotypes.

## ACKNOWLEDGEMENTS

This work was supported by a grant from the NIH (U01DK089569). Human pancreatic islets were provided by the NIDDK-funded Integrated Islet Distribution Program (IIDP) at City of Hope, NIH Grant #2UC4DK098085 as well as the Stanford Islet Research Core. Work in M.H.’s laboratory was supported by NIH R01 (DK105831); Y. K. was supported by a Kraft Family Postdoctoral Fellowship. We are deeply indebted to the islet donors and their families for making this study possible. We are grateful to Dirk Grimm and Stephanie Grosse (University of Heidelberg, Germany) for supplying this study with the AAV backbone vector and the majority of parental capsid sequences. We thank Javier Alcudia (Stanford vector core) for supplying us with several parental capsids and Hiroyuki Nakai (OHSU) for advice on BC library generation. We are indebted to Yan Hang, Heshan Peiris, and Robert Whitener (Stanford University) for help with islet culture, staining and FACS. We thank Hak Kyun Kim (Stanford University) for help with figure generation, Francesco Puzzo for guidance on statistics, and Paul Valdmanis (U. Washington) for development of the conservation and enrichment analysis. This project was also supported by a NIH Shared Instrumentation Grant (S10-OD010580) from the National Center for Research Resources (NCRR) with significant contribution from Stanford’s Beckman Center as well as the Stanford Small Animal Imaging Facility. The authors wish to acknowledge the Stanford Genomics Facility for performing high-throughput sequencing. The contents of this publication are solely the responsibility of the authors and do not necessarily represent the official views of the various funding bodies or universities involved. Packaging plasmids for any of the new capsids described herein must be obtained through an MTA with Stanford University.

## Author contributions

K.P., G.D.A., and M.A.K. designed the experiments. K.P., G.D.A., F.G., M.R.T., Y.K., F.Z., R.S., J.X., S.N., Q.L., J.L., M.H., M.G., and M.A.K. generated reagents, protocols, performed experiments and analyzed data. K.P and M.A.K. wrote the manuscript and generated the figures. All authors reviewed, edited and commented on the manuscript.

