## Supplemental Information for "Using a barcoded AAV capsid library to select for novel clinically relevant gene therapy vectors"

Fig. S1

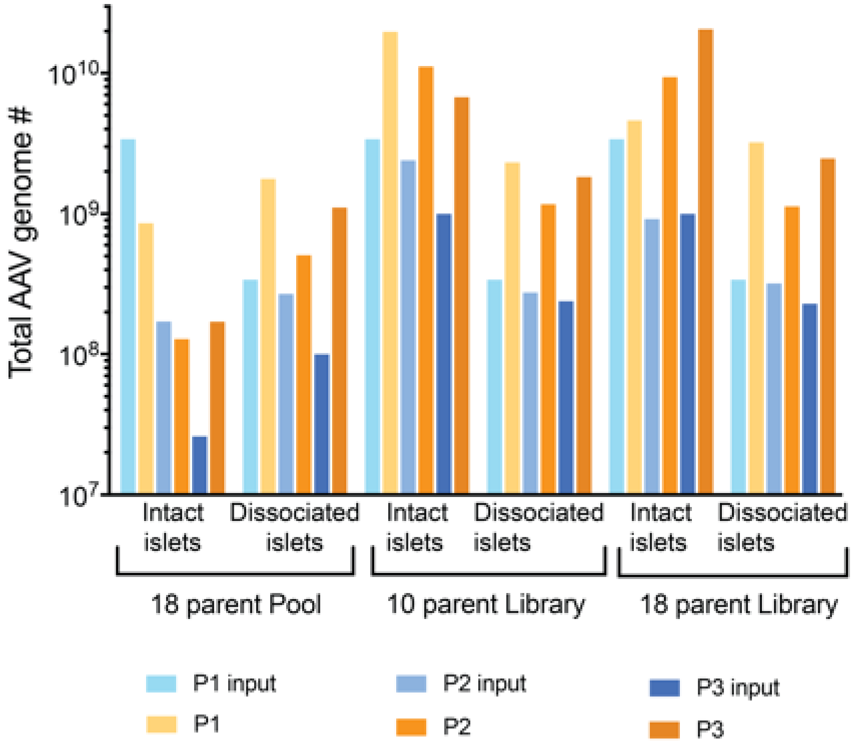


**Figure S1: Replication of the 18 parent pool, the 10 parent library, and the 18 parent library during three rounds of passaging on intact and dissociated islets.** Intact islets were infected with the indicated AAV preparations using an MOI of 20K, dissociated islet cells were infected using an MOI of 2K. Wildtype adenovirus 5 was used to replicate AAV and virus was harvested from the supernatant and cells after 4 days. Replication of viral species was determined by rep qPCR. For each round the total viral genome copies used for infection and the total viral genome copies retrieved are shown.

Fig. S2A

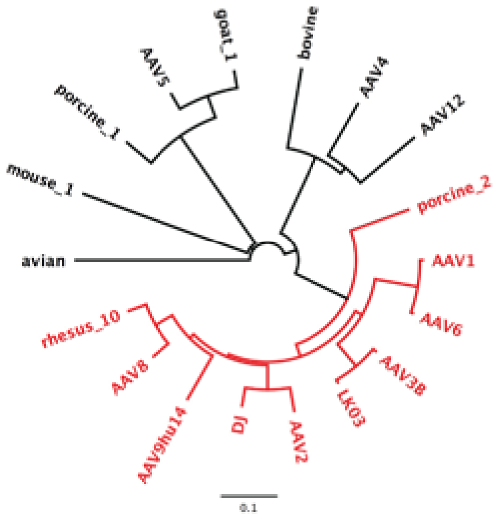


Fig. S2B

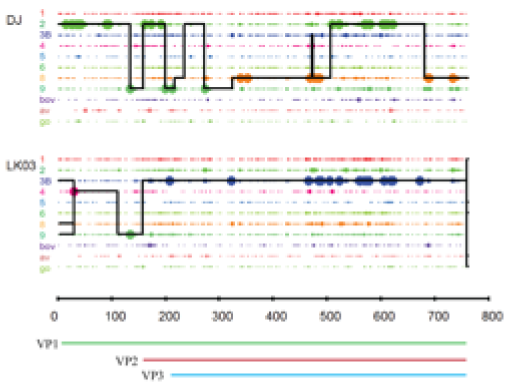


Fig. S2C

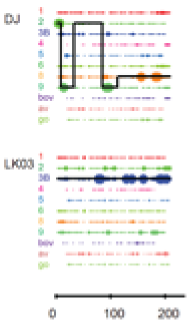


**Figure S2: Parental capsids used for library generation. (A)** Phylogenetic relationship of the 18 parental capsids on the amino acid level. The parental capsids used for generation of the 10 parent library are shown in red. The neighbor-joining tree was constructed using Genious 6.0.6. **(B)** Crossover analysis of chimeric capsids DJ and LK03 with parental contributions indicated. Large dots represent 100% parental match (i.e. the position in question matches only one parent) and small dots represent more than one parental match (i.e. the position matches more than one parent) at each position. The solid line for each chimera represents the library parents identified within the sequence between crossovers. For simple visualization, the same parental capsid sequences are shown for both chimeras although they were derived from different libraries with different parental compositions. AAV-DJ was derived from a library that contained AAV2, AAV4, AAV5, AAV8, AAV9hu14, AAV-bovine, AAV-avian, and AAV-goat1 capsid sequences, AAV- LK03 was derived from a library that contained AAV1, AAV2, AAV3B, AAV4, AAV5, AAV6, AAV8, AAV9hu14, AAV-bovine, AAV-avian, and AAV-goat1 sequences. **(C)** Crossover analysis of AAP for AAV-DJ and AAV- LK03 with parental contributions indicated.

Fig. S3A

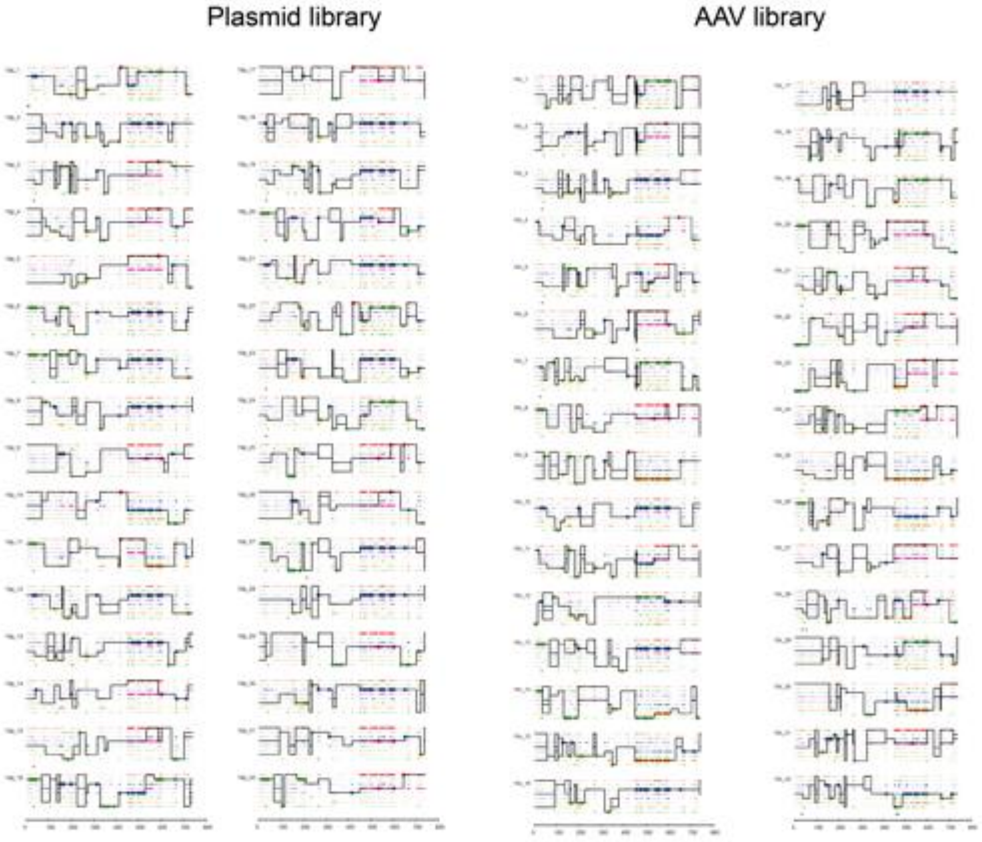


Fig. S3B

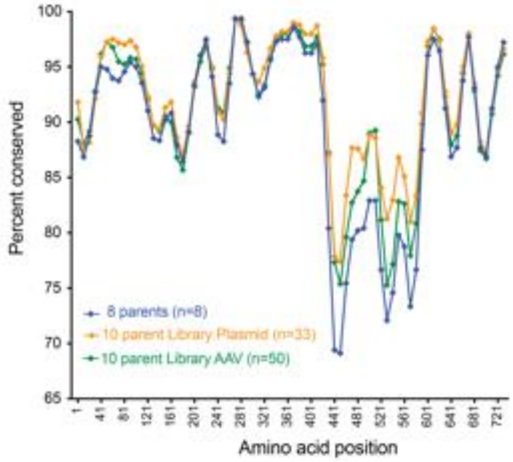


**Figure S3: Analysis of the 10 parent library by Sanger sequencing of random clones. (A)** Crossover analysis of amino acid sequences of several shuffled capsids derived from the plasmid library (left) as well as the AAV library (right). For consistency LK03 and DJ are not shown as individual parents. Positions that are marked with a cross are not *de novo* mutations but were derived from AAV-LK03 that contains a short stretch from AAV4. Parental capsids are shown in the following order: AAV1, AAV2, AAV3B, AAV6, AAV8, AAV9hu14, AAV-rhesus10, AAV-porcine2. **(B)** Conservation analysis along the capsid sequences for the 8 input parental sequences and the 10 parent plasmid as well as rAAV libraries.

Fig. S4A

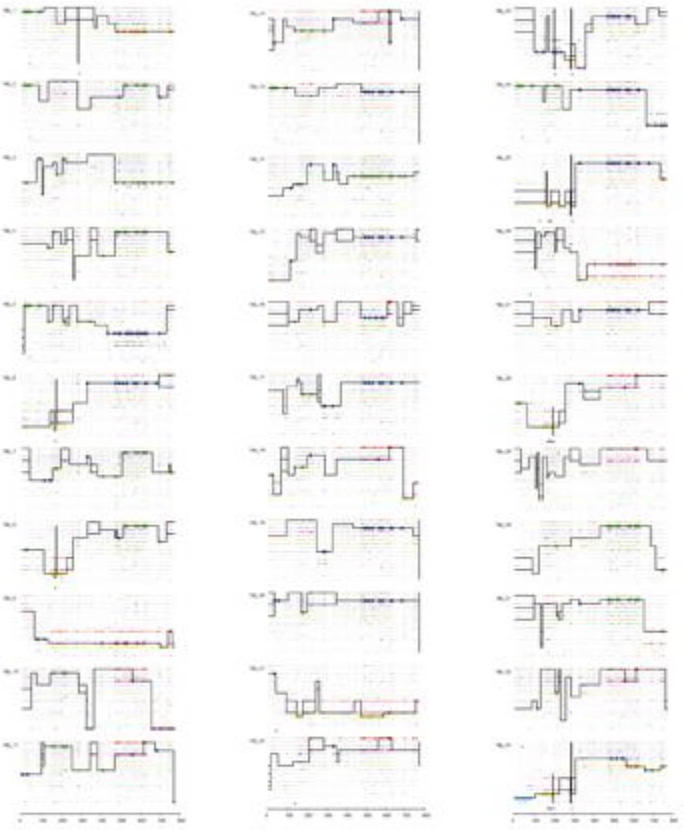

Fig. S4B

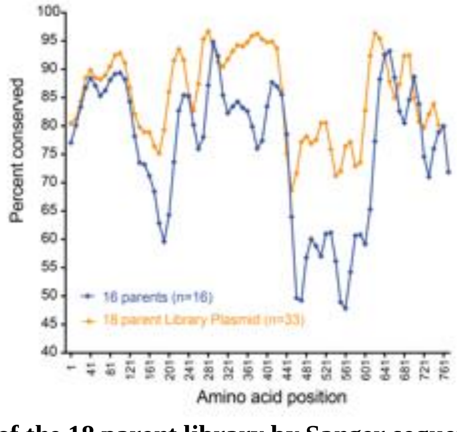


**Figure S4: Analysis of the 18 parent library by Sanger sequencing of random clones. (A)** Crossover analysis of amino acid sequences of several shuffled capsids derived from the plasmid library. Parental capsids are shown in the following order: AAV1, AAV2, AAV3B, AAV6, AAV8, AAV9hu14, AAV-rhesus10, AAV-porcine2, AAV4, AAV5, AAV12, AAV-bovine, AAV-goat1, AAV-porcine1, AAV-mouse1, and AAV-avian. **(B)** Conservation analysis along the capsid sequences for the input 16 parental sequences and the 18 parent plasmid library. Capsids DJ and LK03 are not listed as parents since they are chimeras consisting of fragments from the parentals used.

Fig. S5A

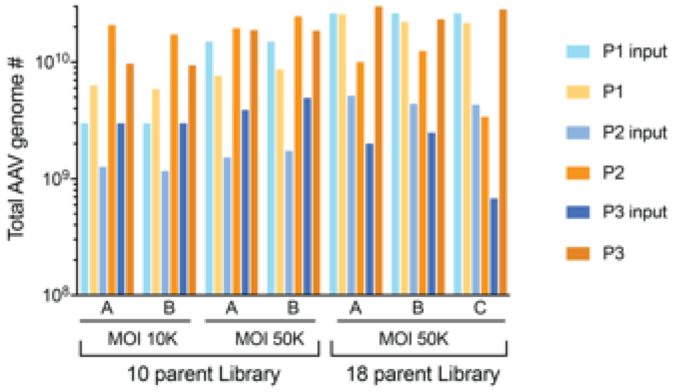


Fig. S5B

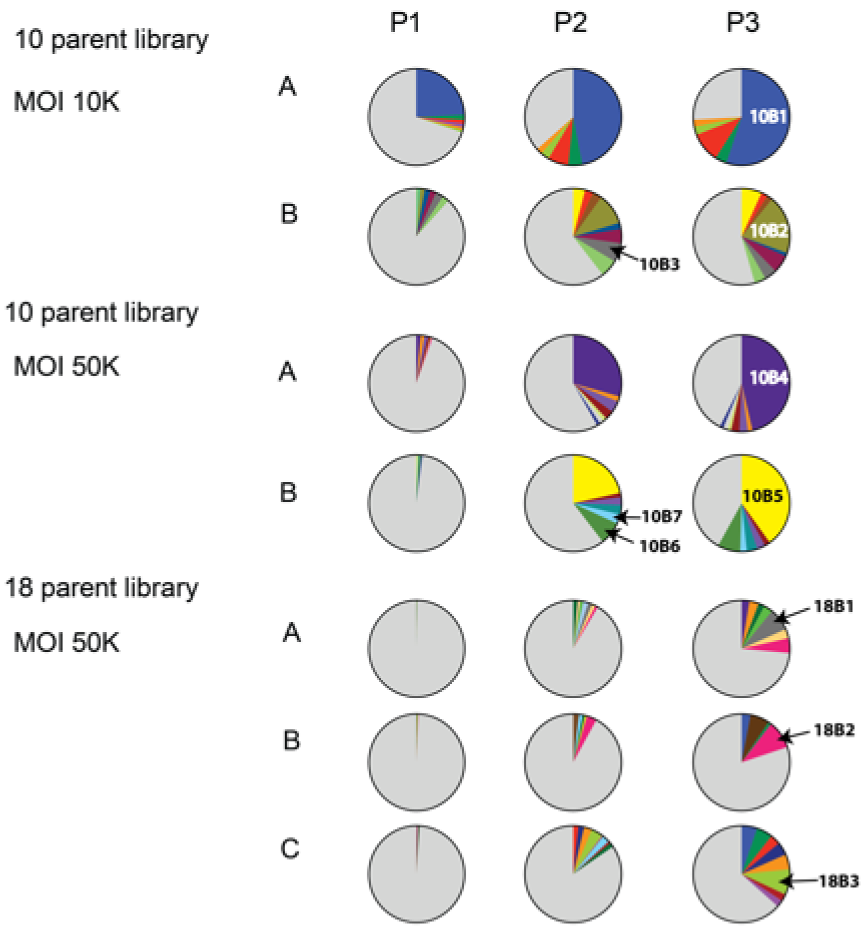


**Figure S5: Results of the second library screen on human islets. (A)** Replication of the 10 parent library and the 18 parent library during three rounds of passaging on intact islets. Islets were infected in biological duplicates with the 10 parent library using an MOI of 10K or 50K. Infection with the 18 parent library was performed in biological triplicates using an MOI of 50K. Adenovirus 5 was used to replicate AAV and virus was harvested from the supernatant and cells after 4 days. Replication of viral species was determined by rep qPCR. For each round the total viral genome copies used for infection and the total viral genome copies retrieved are shown. **(B)** Enrichment of distinct capsids during three rounds of passaging. Barcode sequences were amplified from viral genomes after each passage and were analyzed by high-throughput sequencing. Enriched variants are depicted in different colors while all other variants are shown in grey. Enrichment of AAV capsid variants used for vectorization is indicated (10B1, 10B2, 10B3, 10B4 10B5, 10B6, 10B7, 18B1, 18B2, 18B3).

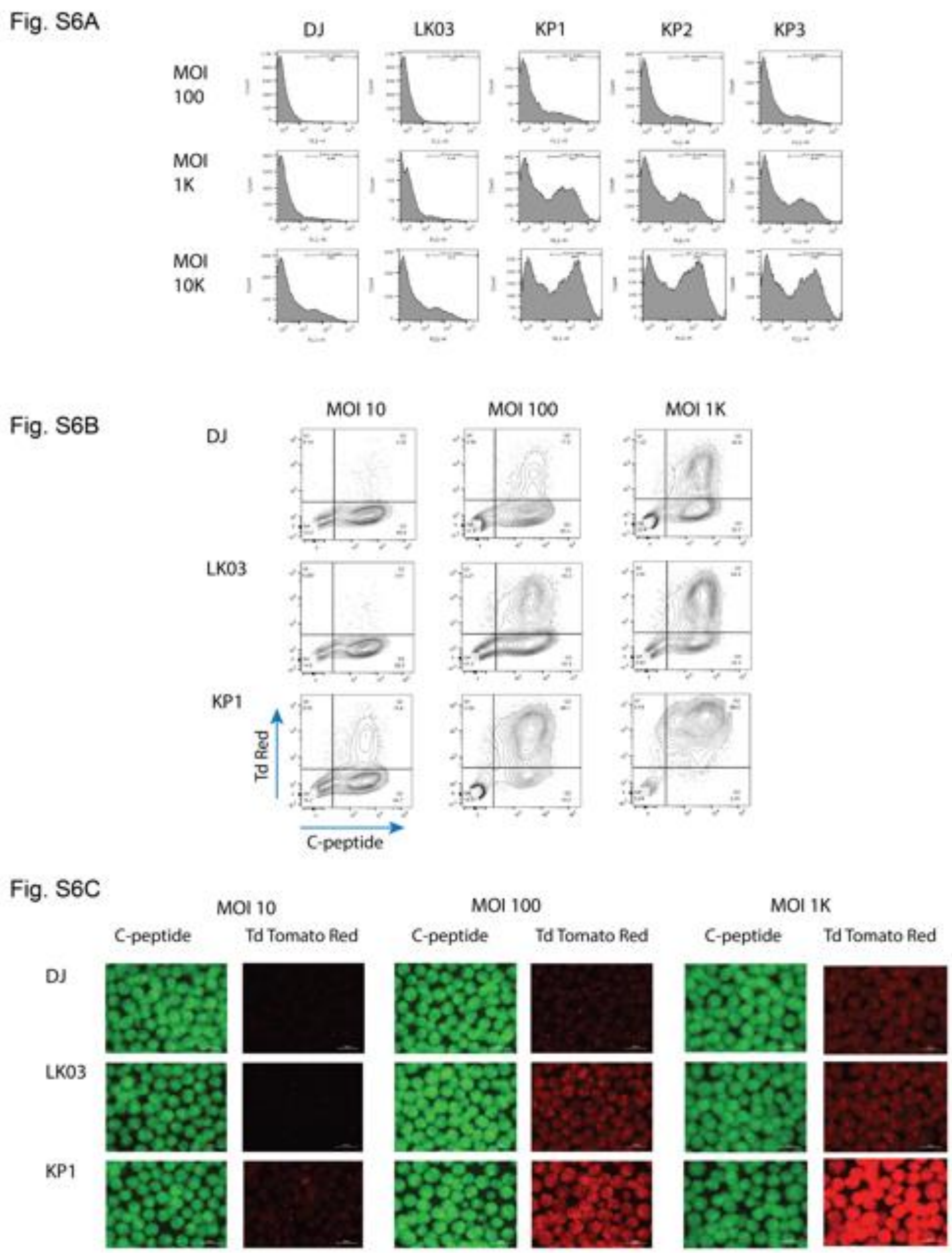


**Figure S6: Transduction efficiency of novel and parental AAV capsids on human islets and on hESC derived β-cells. (A)** Dissociated islet cells were transduced with CsCl gradient purified scCAG-GFP rAAV preps generated with the two best parental capsids as well as the novel capsids that were the top transducers in the pre-screen. Three different MOIs were used for transduction. Transduction efficiency is measured as a function of the number of GFP expressing cells in conjunction with fluorescence intensity for each cell. **(B)** DJ, LK03 and KP1 capsids were used to package a Tomato Red vector and hESC derived mature β-cells were transduced with the MOIs indicated. Intracellular staining for the β-cell marker C-peptide was performed on day 6 post transduction and cells were analyzed by flow cytometry. **(C)** The same cells were also analyzed by immunofluorescent imaging. The size bar represents 200 um.

Fig. S7

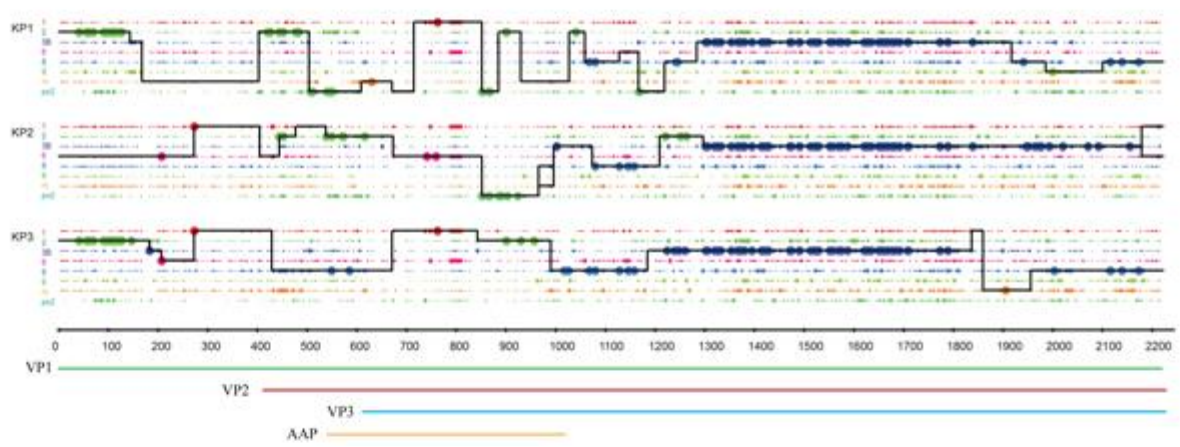


**Figure S7: Analysis of the novel capsids for parental contribution on the nucleotide level.** VP1, VP2, VP3 and AAP coding sequences are indicated below the crossover analysis for KP1, KP2, and KP3 sequences.

Fig. S8A

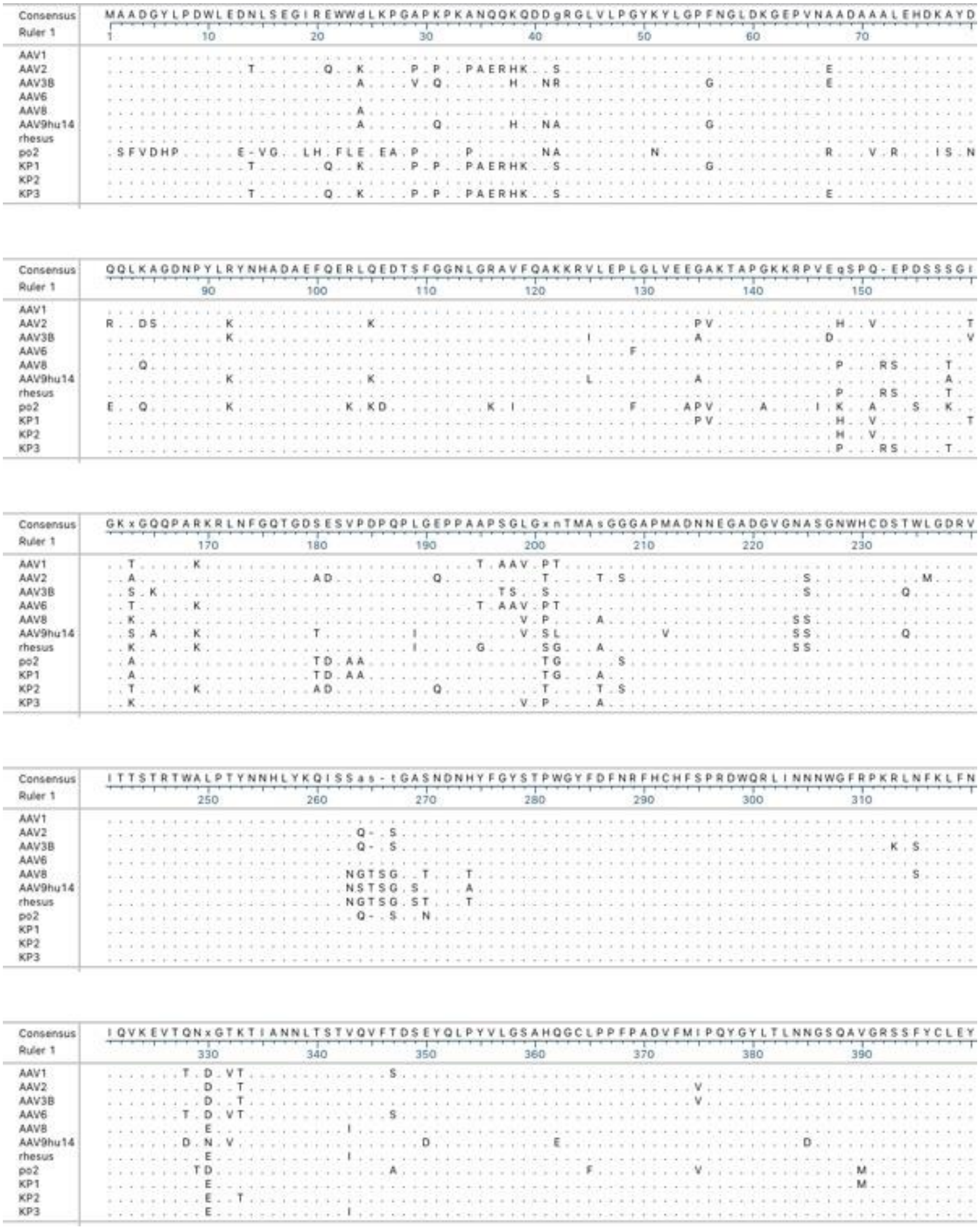
Consensus
MAADGYLPDWLEDNLSEGIREWWdLKPGAPKPKANQQKQDDgRGLVLPGYKYLGPFNGLDKGEPVNAADAAALEHDKAYD
QQLKAGDNPYLRYNHADAEFQERLQEDTSFGGNLGRAVFQAKKRVLEPLGLVEEGAKTAPGKKRPVEqSPQ-EPDSSSGI
GKxGQQPARKRLNFGQTGDSESVPDPQPLGEPPAAPSGLGxnTMAsGGGAPMADNNEGADGVGNASGNWHCDSTWLGDRV
ITTSTRTWALPTYNNHLYKQISSas-tGASNDNHYFGYSTPWGYFDFNRFHCHFSPRDWQRLINNNWGFRPKRLNFKLFN
IQVKEVTQNxGTKTIANNLTSTVQVFTDSEYQLPYVLGSAHQGCLPPFPADVFMIPQYGYLTLNNGSQAVGRSSFYCLEY
Ruler 1
AAV1
AAV2
AAV3B
AAV6
AAV8
AAV9hu14
rhesus
po2
KP1
KP2
KP3

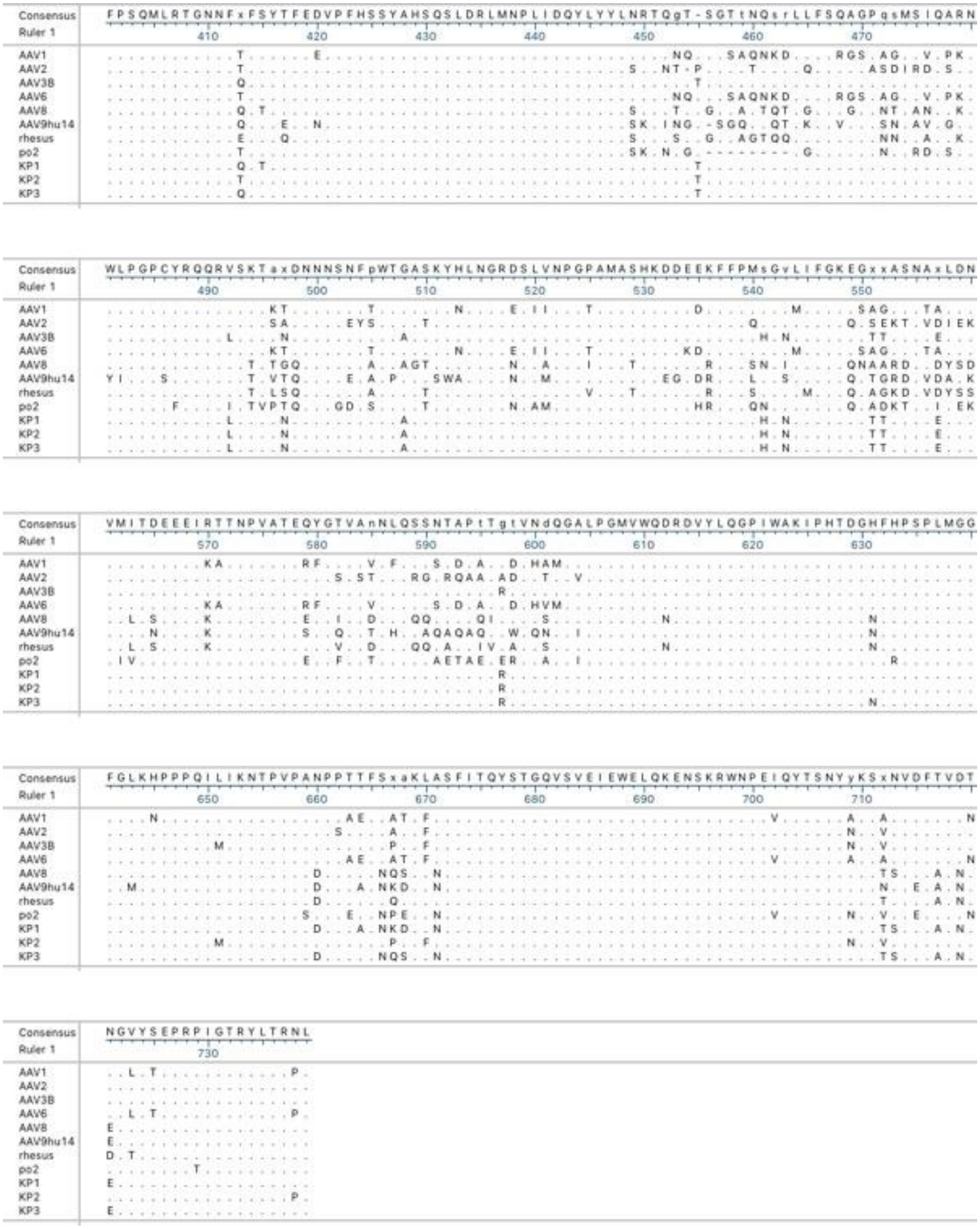
Consensus
Ruler 1
FPSQMLRTGNNFxFSYTFEDVPFHSSYAHSQSLDRLMNPLIDQYLYYLNRTQgT-SGTtNQsrLLFSQAGPqsMSIQARN
410 420 430 440 450 460 470
WLPGPCYRQQRVSKTaxDNNNSNFpWTGASKYHLNGRDSLVNPGPAMASHKDDEEKFFPMsGvLIFGKEGxxASNAxLDN
490 500 510 520 530 540 550
VMITDEEEIRTTNPVATEQYGTVAnNLQSSNTAPtTgtVNdQGALPGMVWQDRDVYLQGPIWAKIPHTDGHFHPSPLMGG
570 580 590 600 610 620 630
FGLKHPPPQILIKNTPVPANPPTTFSxaKLASFITQYSTGQVSVEIEWELQKENSKRWNPEIQYTSNYyKSxNVDFTVDT
650 660 670 680 690 700 710
NGVYSEPRPIGTRYLTRNL
730
AAV1
AAV2
AAV3B
AAV6
AAV8
AAV9hu14
rhesus
po2
KP1
KP2
KP3

Fig. S8B

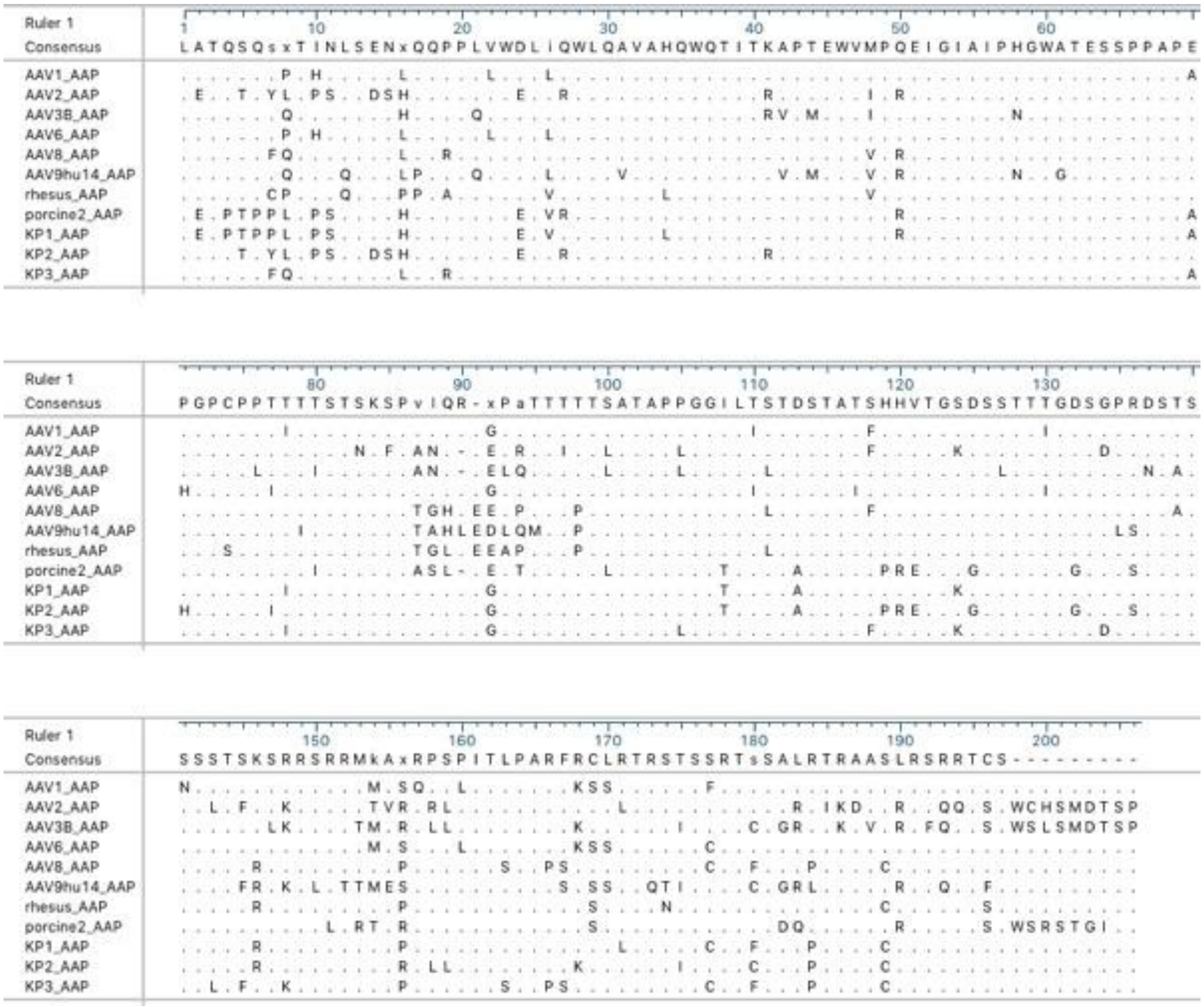


**Figure S8: Amino acid alignments of the novel sequences. (A)** Capsid sequences of the novel variants as well as the 8 parental serotypes were aligned using MegAlign. Only differences to the consensus sequence are shown. The start sites for VP1, VP2, and VP3 are indicated. **(B)** AAP sequences of the novel variants as well as the 8 parental serotypes were aligned.

Fig. S9A

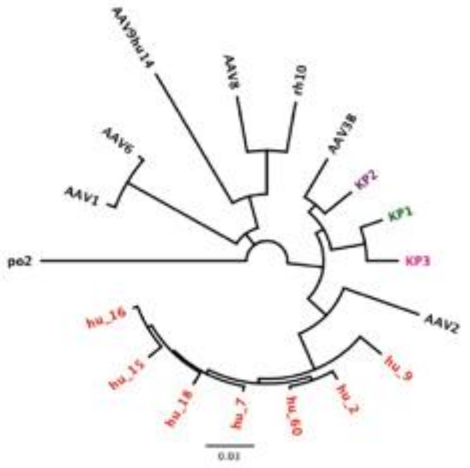


Fig. S9B

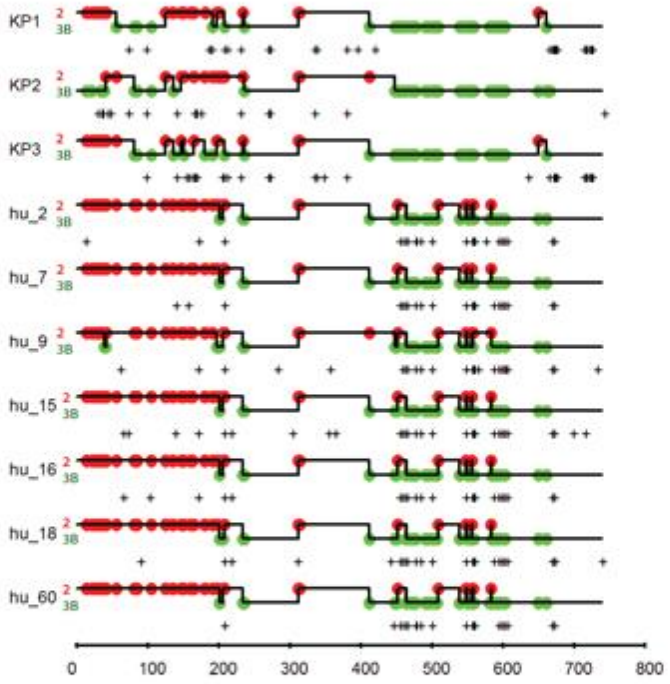


**Figure S9: Parental capsids used for library generation. (A)** Phylogenetic relationship of the 8 capsids used for shuffling as well as the novel capsid variants and several representative capsid sequences from clade C isolates (highlighted in red). The analysis was performed on the amino acid level. The neighbor-joining tree was constructed using Genious 6.0.6. **(B)** Crossover analysis of capsids KP1, KP2, and KP3 as well as clade C capsids using only AAV2 and AAV3B as parentals. Large dots represent 100% parental match (i.e. the position in question matches only one parent) and small dots represent more than one parental match (i.e. the position matches more than one parent) at each position. The solid line for each chimera represents the library parents identified within the sequence between crossovers. A set of thin horizontal parallel lines between crossovers indicates multiple parents match at an equal probability. A mutation is recorded as a plus sign when a position of a chimera does not match the corresponding position in any parental sequence.

Fig. S10A

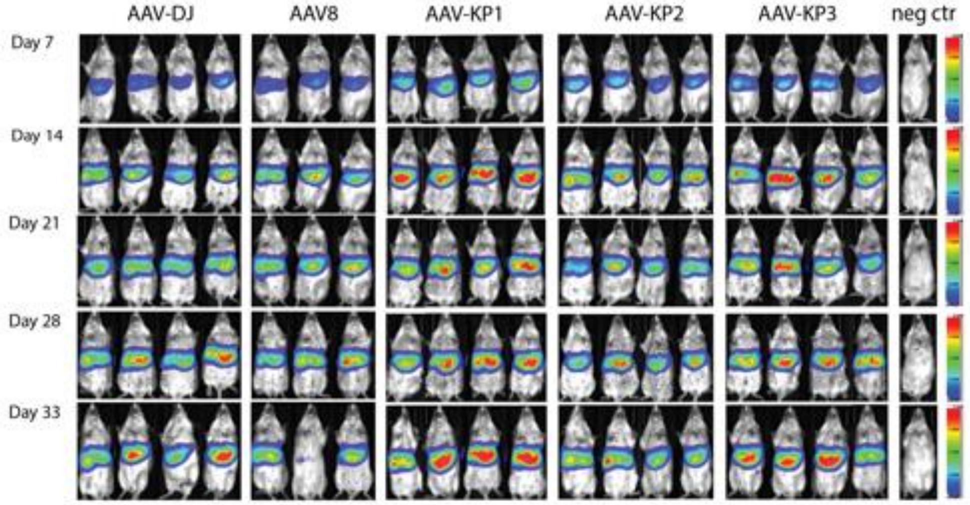


Fig. S10B

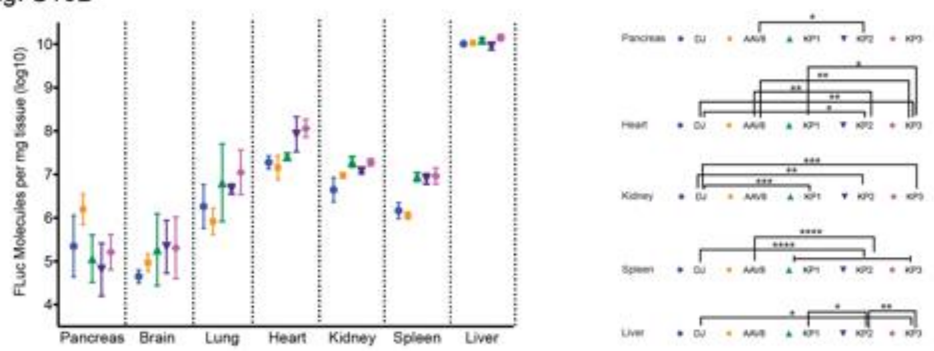


Fig. S10C

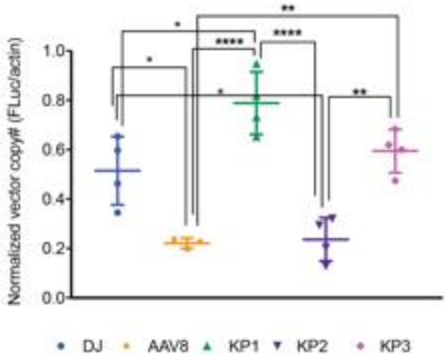


**Figure S10: *In vivo* transduction efficiency of rAAVs packaged with the novel, AAV8, and DJ capsids. (A)** Balb/C SCID mice were injected via tail vein with 2E10 vg of each Firefly luciferase expressing rAAV and luciferase expression in the livers was monitored over several weeks using live imaging after i.p injection of luciferin substrate. Four animals were injected for each group with the exception of the AAV8 group which contained 3 animals. Maximum radiance was set at 2E09 with a threshold of 5E07. **(B)** 34 days post injection mice were sacrificed, and several organs were analyzed for luciferase expression *ex vivo* using the Promega Luciferase assay system. **(C)** Genomic DNA was isolated from mouse livers and analyzed for vector copy numbers by qPCR. Only statistically significant differences between groups are indicated.

*: $p<0.05$, **: $p<0.01$, ***: $p<0.001$, ****: $p<0.0001$

Table S1: Composition of the 18 parental pool based on Illumina HTS of the barcode sequences. Data are given as percentage of total reads with absolute read numbers in parenthesis.

| | | | | | |
|---|---|---|---|---|---|
| AAV1 | 6.65 (68,241) | AAV8 | 5.75 (59,023) | porcine 1 | 5.43 (55,711) |
| AAV2 | 7.79 (79,931) | AAV9hu14 | 5.08 (52,087) | porcine 2 | 9.77 (100,288) |
| AAV3B | 4.74 (48,660) | AAV12 | 0.32 (3,307) | mouse 1 | 4.94 (50,690) |
| AAV4 | 0.01 (62) | DJ | 7.69 (78,973) | goat 1 | 1.47 (15,103) |
| AAV5 | 9.60 (98,474) | LK03 | 6.05 (62,091) | avian | 9.38 (96,275) |
| AAV6 | 1.01 (10,320) | rhesus 10 | 7.67 (78,743) | bovine | 0.00001 (2) |

Table S2: Barcode NGS data of the BC library, and the capsid shuffled libraries at the plasmid as well as the AAV level. The top identical sequences are shown, the number of times each sequence is represented within the sequence reads is shown in parenthesis.

| Sample | BC library | 10 parent plasmid library | 10 parent AAV library | 18 parent plasmid library | 18 parent AAV library |
|---|---|---|---|---|---|
| Total reads (#) | 1,249,614 | 1,256,634 | 1,141,870 | 1,804,277 | 1,000,494 |
| Unique reads (#) | 1,164,408 | 1,110,174 | 866,601 | 1,598,929 | 798,927 |
| Unique reads (%) | 93.18 | 88.35 | 75.89 | 88.62 | 79.85 |
| Identical BC (# of repeats) | 1 (6x) | 7 (7x) | 1 (287x) | 1 (8x) | 1 (232x) |
| Identical BC (# of repeats) | 20 (5x) | 30 (6x) | 1 (195x) | 8 (7x) | 1 (177x) |
| Identical BC (# of repeats) | 342 (4x) | 220 (5x) | 1 (182x) | 63 (6x) | 1 (173x) |
| Identical BC (# of repeats) | 4,844 (3x) | 1,809 (4x) | 1 (177x) | 365 (5x) | 1 (141x) |

Table S3: Shared amino acid residues of the novel variants. Only residues that are different from AAV3B and are shared among at least two of the variants are shown. Amino acids shared among all three variant capsids are highlighted in bold. A blank field indicates a residue identical to that of AAV3B. A missing amino acid is indicated (-). Numbering is according to the alignment in Figure S10A. Hypervariable regions (HVR) in VP3 are labeled according to[89]. Surface exposed residues on the VP3 capsid protein are marked (*).

| Position | Protein / HVR | AAV3B | KP1 | KP2 | KP3 |
|---|---|---|---|---|---|
| 14 | VP1 | N | T | | T |
| 21 | VP1 | E | Q | | Q |
| 24 | VP1 | A | K | D | K |
| 29 | VP1 | V | P | A | P |
| 31 | VP1 | Q | P | K | P |
| 34 | VP1 | A | P | | P |
| 35 | VP1 | N | A | | A |
| 36 | VP1 | Q | E | | E |
| 37 | VP1 | Q | R | | R |
| 39 | VP1 | Q | K | | K |
| **41** | VP1 | N | **D** | **D** | **D** |
| 42 | VP1 | R | S | G | S |
| 56 | VP1 | G | | F | F |
| 67 | VP1 | E | A | A | |
| **92** | VP1 | K | **R** | **R** | **R** |
| **125** | VP1 | I | **V** | **V** | **V** |
| 135 | VP1 | A | P | G | G |
| **147** | VP1, VP2 | D | **E** | **E** | **E** |
| 148 | VP1, VP2 | Q | H | H | P |
| 151 | VP1, VP2 | Q | V | V | |
| 160 | VP1, VP2 | V | T | I | I |
| **165** | VP1, VP2 | K | **Q** | **Q** | **Q** |
| 181 | VP1, VP2 | E | D | D | |
| **197** | VP1, VP2 | T | **S** | **S** | **S** |
| **198** | VP1, VP2 | S | **G** | **G** | **G** |
| 201 | VP1, VP2 | S | T | T | P |
| 206 | VP1, VP2, VP3 | S | A | T | A |
| **225** | VP1, VP2, VP3 | S | **A** | **A** | **A** |
| **234** | VP1, VP2, VP3 | Q | **T** | **T** | **T** |
| **264*** | VP1, VP2, VP3 (HVR I) | Q | **A** | **A** | **A** |
| **266*** | VP1, VP2, VP3 (HVR I) | - | **T** | **T** | **T** |
| **313** | VP1, VP2, VP3 | K | **R** | **R** | **R** |
| 315 | VP1, VP2, VP3 | S | **N** | **N** | **N** |
| **330*** | VP1, VP2, VP3 (HVR II) | D | **E** | **E** | **E** |
| 333* | VP1, VP2, VP3 (HVR II) | T | K | | K |
| **375** | VP1, VP2, VP3 | V | **I** | **I** | **I** |
| 651 | VP1, VP2, VP3 | M | L | | L |
| 660* | VP1, VP2, VP3 | N | D | | D |
| 666* | VP1, VP2, VP3 | S | N | | N |
| 670* | VP1, VP2, VP3 | F | L | | L |
| 671* | VP1, VP2, VP3 | A | N | | N |
| 709* | VP1, VP2, VP3 (HVR IX) | N | Y | | Y |
| 712* | VP1, VP2, VP3 (HVR IX) | V | T | | T |
| 713* | VP1, VP2, VP3 (HVR IX) | N | S | | S |
| 717* | VP1, VP2, VP3 (HVR IX) | T | A | | A |
| 719* | VP1, VP2, VP3 (HVR IX) | D | N | | N |

| | | | | | |
|---|---|---|---|---|---|
| 721* | VP1, VP2, VP3 (HVR IX) | N | E | | E |
